# Efficient profiling of total RNA in single cells with STORM-seq

**DOI:** 10.1101/2022.03.14.484332

**Authors:** Benjamin K. Johnson, Mary F. Majewski, H. Josh Jang, Rebecca A. Siwicki, Marc Wegener, Ayush Semwal, Jacob Morrison, Kristin Gallik, Michael Hagemann-Jensen, Kelly Foy, Larissa L. Rossell, Emily J. Siegwald, Dave W. Chesla, Jose M. Teixeira, Marie Adams, Ronny Drapkin, Corinne Esquibel, Rachael T. C. Sheridan, Ting Wang, Rickard Sandberg, Timothy J. Triche, Hui Shen

**Author notes:** Correspondence should be directed to: Hui Shen Van Andel Institute 333 Bostwick Ave. NE Grand Rapids, MI 49503. These authors contributed equally (Benjamin K. Johnson, Mary F. Majewski). Co-senior authors (Timothy J. Triche, Hui Shen).

## Abstract

Despite significant advances, current single-cell RNA sequencing (scRNA-seq) technologies often struggle with accurately detecting non-coding transcripts, achieving full-length RNA coverage, and/or resolving transcript-level complexity. Many are also difficult to implement or inaccessible without specialized liquid handlers, further limiting their utility. We present Single-cell TOtal RNA-seq Miniaturized (STORM-seq), a random-hexamer primed, ribo-reduced single-cell total RNA sequencing (sc-total-RNA-seq) protocol using standard laboratory equipment. Adapted as a kit, STORM-seq constructs sequence-ready libraries in one working day, producing the highest complexity scRNA-seq libraries to-date, robustly measuring transcript isoforms and clinically relevant gene fusions in single cells. STORM-seq faithfully reconstructs expression profiles of locus-level transposable elements (TEs), and provides high-resolution profiling of transient, low-abundance enhancer RNAs (eRNAs), offering a powerful tool to dissect single-cell gene regulatory networks in unprecedented detail. Applied to human fallopian tube epithelium, the improved transcriptional resolution reveals a putative progenitor-like population and intermediate cell states, shaped by TEs and non-coding RNAs.

## Introduction

Over the past decade, single-cell sequencing technologies have been developed to assess the heterogeneity of cells across organisms, tissues, cell types and states^1^. Whole-organism and tissue-level single-cell atlases have been generated using a variety of approaches^2,3^, with the most common being single-cell RNA-seq (scRNA-seq). This has spurred rapid development of both new library preparation techniques and associated computational tooling^4^. Recent advances in plate-based scRNA-seq have been developed to assess single-cell whole transcriptomes at high resolution – exemplified by Smart-seq3xpress (SS3x), VASA-seq, Smart-seq-total, and others^5–9^. Profiling polyadenylated and non-polyadenylated transcripts (total RNA) in single cells (sc-total-RNA-seq) has greatly expanded our understanding of transcriptional regulation and coordinated gene expression changes in cell states and fates^5–9^. Generation of scRNA-seq libraries has largely coalesced to a shared set of steps: isolation of single cells (e.g. microfluidics or flow cytometry), cell lysis, reverse transcription (RT), addition of unique molecular identifiers (UMIs) and cell barcodes, pooling, enrichment of transcripts of interest (e.g. mRNA or non-rRNA), and sequencing. The implementation of these steps, resulting data density, and mitigation of protocol-specific technical biases have continued to drive the introduction of novel scRNA-seq methods to better address biological questions.

Efforts to profile total RNA in single cells have typically taken the approach of polyadenylating total RNA followed with amplification utilizing an oligo(dT) primer (VASA-seq, Smart-seq-total)^6,7^. This is different from the random-hexamer priming approach commonly used in traditional bulk total RNA. These approaches are taken likely due to the potential for random hexamers to prime and amplify genomic DNA (gDNA) if present during RT^10^. However, it is known that oligo(dT) approaches can prime genomic poly-A tracks if gDNA is present^11,12^. To the best of our knowledge, this potential background in current polyA-based sc-total-RNA-seq methods has not been examined. We reasoned that random hexamer priming would provide the most sensitive sc-total-RNA-seq approach by omitting the long poly-adenylation steps and mitigating potential gDNA contamination during cell lysis, moving quickly from lysed cells to RT. All scRNA-seq methods that use template switching oligos (TSOs) have the potential to introduce chimeric transcripts during RT and second-strand synthesis^13^. Recent scRNA-seq methods capturing polyadenylated RNA have shown that the design and composition of the nucleotide sequence of the UMI-TSO is critical to minimize these artifacts^5,8^. However, existing sc-total-RNA-seq technologies using UMI-TSOs (e.g. VASA-seq, Smart-seq-total, SnapTotal-seq) are unable to readily identify the impact of contaminating strand invasion artifacts due to methodological choices^6,7,9^. As a result, strand invasion artifacts in sc-total-RNA-seq remain poorly characterized, potentially skewing gene expression profiles and ultimately, biological interpretations.

The ability to account for cell-to-cell RNA content differences is an additional layer of bias that can distort expression profiles if left unaccounted for (e.g. *MYC* amplification and cell-cycle)^14,15^. Indeed, the External RNA Controls Consortium RNA spike-in mix (ERCCs)^16^, Spike-in RNA Variant Control mixes (SIRVs), molecular spikes^17^, and others have been developed to accomplish these tasks, enabling the ability to account for RNA content differences during analysis. However, when and how spike-ins are added during library preparation requires thoughtful consideration to limit introduction of technical biases and consumption of valuable sequencing reads^14^.

With the promise of scRNA-seq revealing heterogeneity in single-cells, it is important to consider going beyond gene counts alone. Transposable elements (TE) and TE-derived transcripts are now well-appreciated to play critical roles in species evolution and tissue development through their intrinsic transposition capabilities and co-option, providing a rich set of gene regulatory features such as enhancers and promoters^18–20^. Under stress, cryptic regulatory elements within transposable elements (TEs) undergo epigenetic reactivation in cells, contributing to disease progression — particularly oncogenesis — by promoting oncogene expression through a mechanism known as onco-exaptation^21–23^. However, existing scRNA-seq technologies struggle to reconstruct the expected TE expression profiles in single cells, exhibiting a preferential bias toward the expression of short interspersed nuclear elements (SINEs)/Alu elements over long interspersed nuclear elements (LINEs)/long terminal repeats (LTRs)^24^. Furthermore, the data sparsity issue of current scRNA-seq approaches limits the detection of TE expression heterogeneity within cell populations and tumors. Despite methodological advancements, new single-cell technologies that comprehensively recapitulate expected locus-level TE and TE-derived transcript expression profiles are needed.

Another challenging class of non-coding transcripts to assay in single cells is enhancer RNA (eRNA), due to their short half-lives (1-2 minutes on average) and low expression^25–27^. Specialized assays to identify these transient RNA species have been developed, with nuclear run-on followed by cap-selection assays (GRO/PRO-cap) being the most sensitive in detecting eRNAs^28^. Further, transient transcriptome sequencing (TT-seq) and related single-cell nascent transcript sequencing has enabled capturing short and long-lived RNA species over time to estimate rates of RNA synthesis and decay, including eRNAs^25,26,29,30^. Bulk and scRNA-seq methods have shown that eRNAs can be captured in principle, though to a lesser extent than their specialized counterparts due to primarily profiling steady-state RNA^28^. However, these nascent RNA methods typically require additional steps that are often not compatible with profiling primary tissues^31^. As a result, high sensitivity sc-total-RNA-seq methods to measure eRNAs and their contribution to cell states and fates, are an important gap to fill between specialized approaches and traditional scRNA-seq methods.

Many contemporary scRNA-seq methods are time consuming to implement and execute, and challenging to analyze, increasing the barrier to entry for these methods. To address these barriers, we present STORM-seq, a single cell total RNA sequencing method that has been adapted as a commercially available kit, makes use of random hexamer priming, incorporation of unique molecular identifiers (UMI), requires no specialized equipment for library preparation, leverages spike-in controls, and generates the most complex scRNA-seq libraries to date. We compare STORM-seq and current, sc-total-RNA-seq protocols in HEK293T and K-562 cell lines, demonstrating STORM-seq more robustly and sensitively profiles total RNA. Further, we show that this method can reconstruct transposable element (TE) expression profiles in single cells, capture short-lived enhancer RNAs (eRNAs), and clinically relevant gene fusions. We apply STORM-seq to the human pre-menopausal fallopian tube epithelium (FTE). FTE plays important roles for reproductive biology, and likely harbors the cell of origin for high-grade serous ovarian cancer. However, the stem/precursor population for this important tissue has been elusive and prior single-cell studies yielded divergent results^32–34^. With the increased resolution of STORM-seq, we discover a putative progenitor-like population and intermediate cell types, with potential long intergenic non-coding RNAs (lincRNAs) and TEs as candidate drivers shaping lineage fate in this tissue.

## Results

### Development of STORM-seq

STORM-seq is built to allow use of off-the-shelf reagents and standard equipment, addressing various limitations of other methods (Supplemental Table 1). Single cells are index sorted into a microwell plate containing Fragmentation Buffer (FB) so that the flow cytometric phenotype of each sorted cell will be associated with the well name for downstream data integration. After cells are lysed, RNA is fragmented for 3 minutes. This optimized fragmentation time generates longer RNA fragments enabling longer paired-end sequencing, allowing for more even coverage across single cells (**Supp.** Fig. 1a-d)^35^. Additionally, fragmenting RNA immediately versus waiting to fragment after conversion to cDNA has been shown to reduce random priming bias^35,36^. By utilizing random priming, STORM-seq captures total RNA without additional polyadenylation steps. Following the random priming of total RNA, STORM-seq utilizes an MMLV-derived reverse transcriptase (RT) with template switching (TS) functionality to add a unique molecular identifier (UMI) sequence during first strand cDNA synthesis. ERCC spike-in controls are also added at the RT and TS step. Introducing ERCCs at this stage prevents fragmentation of the ERCC transcripts, ensuring that counts are accurate after UMI collapsing^15^. Next, PCR amplification is performed, generating double stranded cDNA libraries, each containing a unique dual index (UDI) for multiplexing single cells. At this stage, all wells of the plate are combined into a single pool^37^. With a larger, pooled volume, the remainder of the protocol mimics a bulk total RNA-seq library preparation and can be performed with a single channel pipette. Alternatively, pooled plates can be frozen and stored, allowing multiple plates to be processed through the rest of the protocol simultaneously, increasing throughput and decreasing technical plate-to-plate bias. Ribosomal RNA (rRNA) comprises 80-90% of the RNA within a cell and is typically depleted prior to sequencing total RNA^38–40^. STORM-seq uses a probe-based targeted enzymatic digestion to remove rRNA. Depleting the rRNA from a larger pool has the additional benefit of preserving lowly expressed transcripts, likely due to the rRNA acting as a “carrier” to protect the mRNA and ncRNAs through the enzymatic digestion step and subsequent cleanup^41^. The rRNA-depleted pool is then PCR amplified to introduce sequencing adapters, a final cleanup is performed, and sequenced (**Fig. 1a-b**). Following targeted, probe-based rRNA depletion, STORM-seq libraries contained <2% rRNA content in each cell (**Supp.** Fig. 1e). We have also implemented an automated quality control pipeline to collect various metrics post-sequencing of STORM-seq libraries (**Supp.** Fig. 2**, Methods**). This protocol is modular, flexible, and can be pipetted manually or using an automated liquid handler, going from live cells to sequence-ready libraries in one working day – the fastest end-to-end sc-total-RNA-seq protocol (**Fig. 1c**).

**Figure 1.**
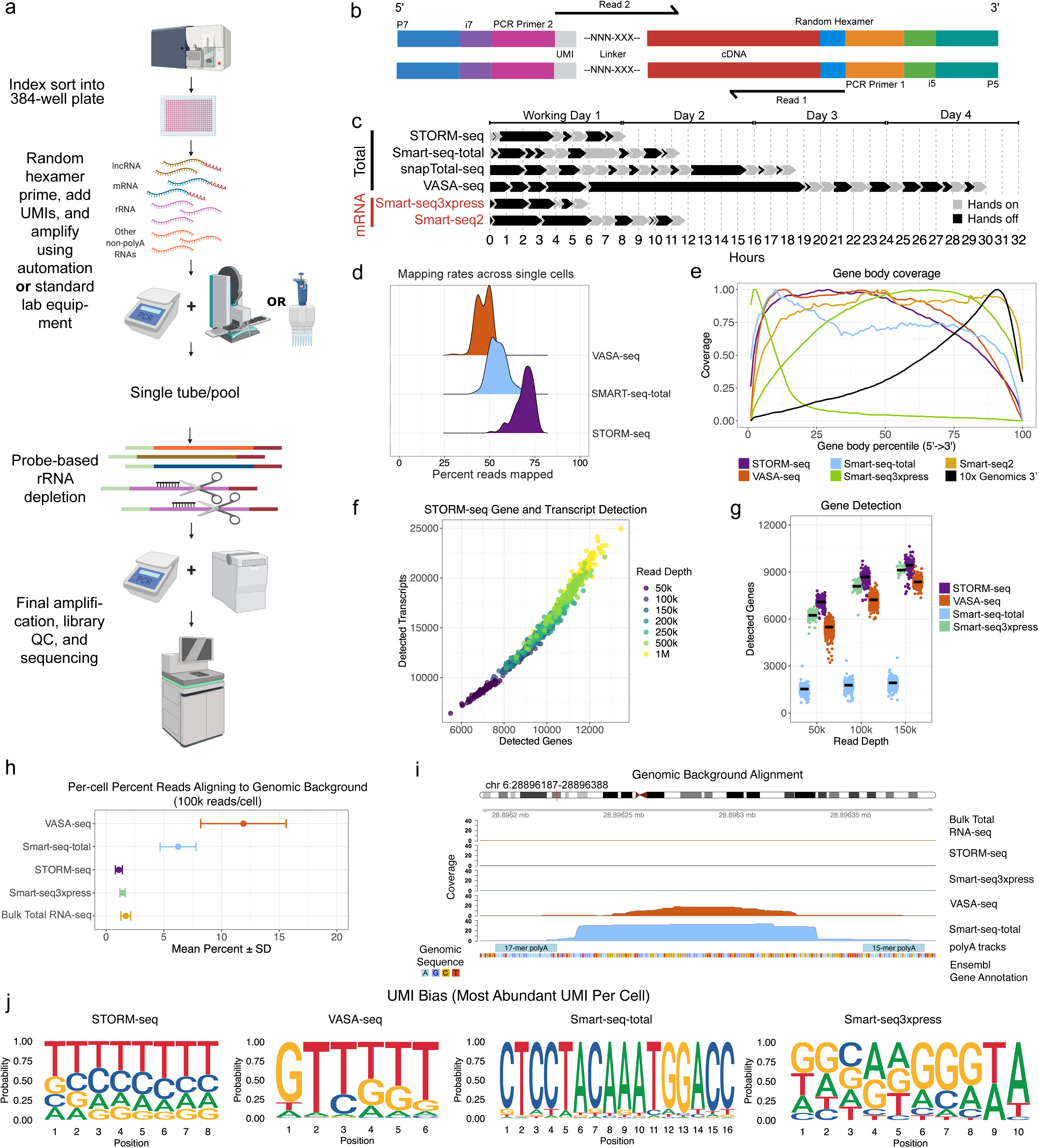
STORM-seq efficiently profiles total RNA in single-cells. **a)** Overview of the STORM-seq library preparation protocol. **b)** Fragment structure and annotations of final STORM-seq libraries. **c)** STORM-seq is the fastest single-cell total RNA-seq method, similar in total time to Smart-seq3xpress. Hands on time: active handling/pipetting of cells/samples; Hands off time: samples/cells not being actively handled (e.g. PCR steps). **d)** STORM-seq has the highest mapping rates in single-cells compared to VASA-seq and Smart-seq-total. **e)** STORM-seq has full gene-body length coverage, similar to other plate-based approaches. Protein coding genes only shown for comparison purposes. **f)** STORM-seq identifies thousands of genes and transcripts per cell across read depths. **g)** Gene detection rate comparison demonstrates STORM-seq measures the most genes/cell compared to other methods (UMI count minimum of 1). **h)** Background genomic alignment percentages in single cells (100k reads/cell) demonstrates that STORM-seq aligns similar proportions of reads to known coding, non-coding, and intergenic TE space, similar to bulk total RNA-seq. **i)** Example genomic background alignment coverage across technologies shows regions with coverage are flanked by poly-A sequences. **j)** Sequence logo plots of the most abundant UMI per cell across technologies. Expected results are even repre- sentation for random UMIs for STORM (NNNNNN - 8 bp UMI), VASA-seq (NNNNNN - 6 bp UMI/UFI), Smart-seq-total (16xN - 16 bp UMI), and Smart-seq3xpress (NNNNNNNNWW - 8bp random + 2 bp W (A/T)). All technology comparisons and results shown are in HEK293T cells at subsampled sequencing depths shown.

### Benchmarking STORM-seq against current, plate-based scRNA-seq protocols

Alignment/mapping, and quantification of scRNA-seq is often performed using two approaches: 1) pseudoalignment to the transcriptome (e.g. kallisto|bustools)^42^ and 2) splice-aware alignment to the genome (e.g. STARsolo)^43^. To facilitate STORM-seq data analysis, the library fragment structure (**Fig. 1b**) exists as a preset within kallisto|bustools. Additionally, we have developed a tool to add synthetic cell barcodes to STORM-seq data called “synthbar” to integrate seamlessly with STARsolo (**Methods**). To the best of our knowledge, STORM-seq is the only paired-end sc-total-RNA-seq method, and when combined with the innovative library preparation approach, we observe more usable reads per cell post-transcriptome alignment, compared to VASA-seq and Smart-seq-total (**Fig. 1d**). Given that STORM-seq is a full-length (gene-body) protocol, we compared gene-body coverage across current scRNA-seq methods. Indeed, we find that STORM- seq covers the gene-body (**Fig. 1e**) and expected gene-length detection bias, similar to other full-length methods (**Supp.** Fig. 3a). STORM-seq produces high-complexity transcript isoform- and gene-level libraries across sequencing depths (**Fig. 1f, Supp.** Fig. 3b-c). Further, STORM-seq recovers more genes per cell compared to the latest sc-total- RNA-seq protocols, VASA-seq and Smart-seq-total, as well as the latest mRNA protocol, Smart-seq3xpress (SS3x; **Fig. 1g**).

Genomic DNA (gDNA) contamination is a primary concern for random-primed total RNA-seq, as MMLV-derived RT will amplify DNA, as well as RNA^10^. Additionally, oligo(dT)- primers in the presence of gDNA will prime DNA containing poly-A tracks, found throughout the genome^11,12^. We reasoned that background genomic alignments may serve as a proxy for spurious priming events if gDNA is present. Therefore, we investigated reads aligning to unannotated coding and non-coding space, combined with known transposable element (TE) annotations, and not found within known R-loop regions. STORM-seq (random primed) and SS3x (oligo(dT) primed) exhibit minimal genomic background alignments, similar to bulk total RNA-seq (random primed). In contrast, VASA-seq (oligo(dT) primed) and Smart-seq-total (oligo(dT) primed) have up to ∼15% genomic background alignments, often found to be flanked by poly-A runs, though may be more prevalent given the read depth and filtering performed (**Fig. 1h-i**, **Methods**).

UMI diversity is critical for mitigating the technical effects of PCR amplification bias, with downstream consequences of under- or over-collapsing UMIs if systematic bias persists during library preparation, sequencing, and data analysis^42^. To examine UMI bias across technologies, we constructed per cell UMI sequence logos to visualize diversity (unique UMIs), prevalence (frequency of UMIs), and bias (most abundant UMI per cell), as well as the observed/expected inter-gene UMI collision rates. Based on the random nucleotide construction for each UMI (except for SS3x which has the added 3’ WW nucleotide motif, **Supp.** Fig. 4d-e), we reasoned that the base diversity at each position in the UMI should be evenly represented, across metrics. Indeed, STORM-seq, VASA-seq, and SS3x had relatively even representation of each base across UMI diversity and prevalence, indicating expected starting UMI diversity during library preparation and carried through to data analysis. In contrast, Smart-seq-total had biased base diversity across the length of the 16bp UMI (**Supp.** Fig. 4d-e). Next, we examined the most abundant UMI sequence per cell and found that STORM-seq has the most even representation of the expected random nucleotide diversity across the length of the UMI, with VASA-seq and Smart-seq-total being the most severely affected (**Fig. 1j**). To estimate inter-gene collision rates, we constructed a simulation framework as a function of UMI length and gene detection (e.g. sequence depth) to establish the expected collision rates across UMI lengths/base diversity in the technologies examined (**Supp.** Fig. 4a, **Methods**). Indeed, UMI inter-gene collision rates decreased as the length of the UMI increased from 6bp (VASA-seq) to 16bp (Smart-seq-total) in the simulation results. We observe the expected UMI inter-gene collision rates in STORM-seq and VASA-seq, with the most severe in Smart-seq-total (∼7x the expected rate; **Supp.** Fig. 4b). This elevated inter-gene collision rate is likely explained by the observed versus expected UMI saturation, with the largest effects being shown when UMI saturation is low (**Supp.** Fig. 4c). Moreover, STORM-seq exhibits low strand invasion artifacts, similar to SS3x (**Supp.** Fig. 4f, **Methods**). Strand invasion artifacts were not able to be calculated for VASA-seq and Smart-seq-total, due to the read architecture.

### Robustness and sensitivity of transcript detection

Typical metrics for new scRNA-seq technologies include quantification of detected genes per cell, library construction and throughput optimizations, library complexity, and applications that may improve biological insights^5–7^. While these metrics help contextualize new technologies within contemporary methods, it is important to assess whether they provide better, more robust measurements. Here we propose that simulation of multiple experiments through subsampling and resampling techniques across technologies and a common cell type (HEK293T), allows demonstration of expected transcript detection rates and transcript expression variance (robustness) in sc- total-RNA-seq methods in single-cells (**Supp.** Fig. 5a, **Methods**). Simulation of 10 experiments through randomly subsampling single HEK293T cells to 50k reads/cell (**Supp.** Fig. 5a), showed that STORM-seq has greater transcript detection rates across annotated transcript lengths, compared to VASA-seq and Smart-seq-total (**Supp.** Fig. 5b). Moreover, STORM-seq produces the most robust (lowest variance) transcript abundance estimates across transcript lengths, compared to VASA-seq and Smart-seq-total (**Supp.** Fig. 5c). Taken together, STORM-seq produces more consistent, robust transcript abundance estimates compared to contemporary sc-total-RNA-seq methods.

Next, we examined the sensitivity of STORM-seq to capture single molecules in individual cells by using ERCC spike-in transcripts that are expected to be at ∼1 copy per cell at the 1:1 million (M) dilution used, across 3 different cell types and sequencing depths (**Methods**)^44^. Indeed, STORM-seq reconstructs expected ERCC spike-in copy number that have at least ∼1 molecule per cell across all cell types (HEK293T, K-562, and RMG- 2) and sequencing depths tested (≥0.9 median adjusted R^2^ for observed versus expected copy number; **Supp.** Fig. 5d-e). Moreover, STORM-seq showed a median sensitivity of single molecule ERCC spike-in detection of ∼88% at 100k reads/cell and increases with sequencing depth, across cell types, making STORM-seq the most sensitive sc-total-RNA-seq method to date (**Supp.** Fig. 5f)^45^. STORM-seq also demonstrates good cell-to-cell gene expression correlations within the cell types tested (**Supp.** Fig. 5g).

### Reconstruction of TE transcripts, regulatory elements, and gene fusions in single-cells

To investigate the capability of STORM-seq to recapitulate bulk total RNA-seq (current “gold-standard”) TE profiles in single-cells, we calculated the observed over expected scores comparing transcribed and genomic TE family representation as previously described^24^. STORM-seq shows the most consistent TE family representation in HEK293T (**Fig. 2a)** and K-562 **(Supp.** Fig. 6a) single-cells, similar to bulk total RNA-seq, across scRNA-seq technologies tested. Long interspersed nuclear elements (LINEs) L1 and L2, and long terminal repeats (LTRs) ERV1 and ERVK contain endogenous promoters that are often exapted in a cell-type specific manner^18^. STORM-seq robustly captures these TE families in single cells compared to other scRNA-seq methods . Short interspersed nuclear elements (SINEs) are over-represented in VASA-seq and Smart-seq-total relative to bulk total RNA-seq, likely due to the oligo(dT) primed library construction strategy of these methods and the preferential genomic location of SINEs found at the 3’ ends of genes (**Fig. 2a, Supp.** Fig. 4a)^46^. Moreover, STORM-seq has the highest locus-level TE expression correlation to bulk total RNA-seq (R^2^ of 0.655) compared to VASA-seq (R^2^ of 0.162) and Smart-seq-total (R^2^ of 0.003; **Fig. 2b**). These results demonstrate that STORM-seq is the only scRNA-seq method that can faithfully reconstruct TE family and locus-level expression profiles in single cells.

**Figure 2.**
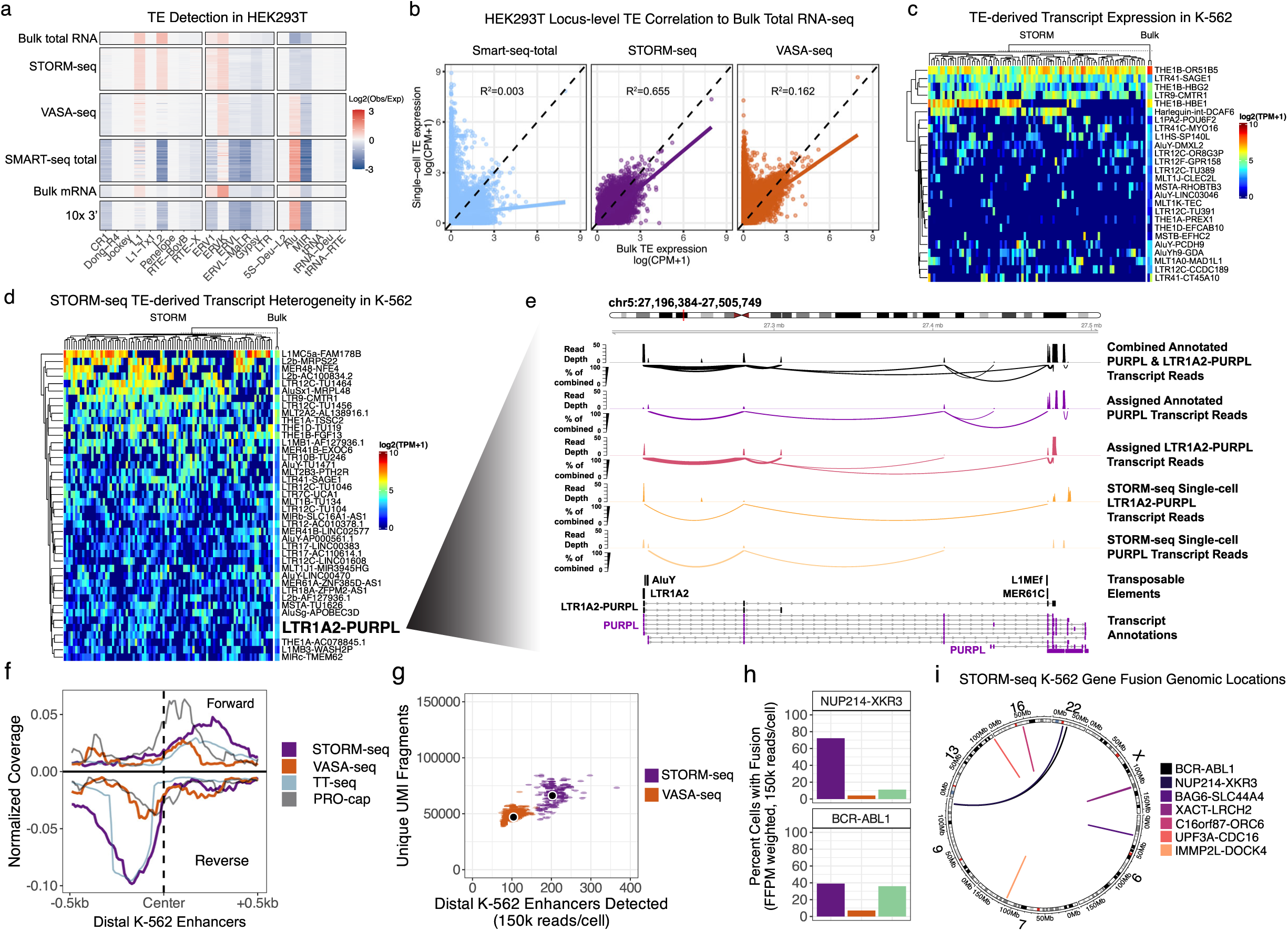
STORM-seq reconstructs cell-type specific regulatory elements and clinically relevant gene fusions in single-cells. a) Sin-gle-cell transposable element (TE) expression representation (obs/exp) across single-cell technologies and bulk RNA-seq. Bulk total RNA-seq is considered the “gold standard”. STORM-seq reconstructs TE profiles in single cells similar to bulk total RNA-seq. **b)** STORM-seq quantifies locus-level TE expression similar to bulk total RNA-seq. **c)** STORM-seq reveals single cell TE-derived transcript expression heterogeneity in previously reported bulk-level TE-derived transcript candidates in K-562 (Shah *et al.,* Nat. Genetics 2023), demonstrating that bulk-level discov- ery of TE-derived transcripts does not necessarily indicate ubiquitous expression across single cells. **d)** Discovery and characterization of addi- tional TE-derived transcript expression heterogeneity found in bulk total RNA-seq. LTR1A2-PURPL is being shown as an example of consistent expression across replicates in bulk total RNA-seq, but heterogeneous expression across K-562 single cells. Transcript names beginning with “TU” are unannotated genes/transcripts found in Ensembl 101 annotations. TE-derived transcripts above and below LTR1A2-PURPL have been removed for aesthetic purposes but the full list can be found in Supplemental Table 3. **e)** STORM-seq reads spanning annotated PURPL tran- scripts and LTR1A2-PURPL TE-derived transcripts are shown in pseudobulk (combined and assigned annotated PURPL and LTR1A2-PURPL tracks) and single cells with associated splice junction spanning reads as a percent of the maximum spanning read depth found in the combined track. The LTR1A2-PURPL TE-derived transcript comprises the majority of PURPL expression as shown by the percentage of total junction spanning reads in the assigned LTR1A2-PURPL track. **f)** Stranded coverage profiles of pseudobulk STORM-seq and VASA-seq across intergen- ic, distal K-562 enhancers, with transient transcriptome sequencing (TT-seq) and PRO-cap coverage. **g)** Detection rates of single-cell distal eRNAs when subsampled to 150k reads/cell shows STORM-seq identifies approximately twice as many eRNAs per cell compared to VASA-seq. **h)** Proportion of single cells with a detected, known gene fusion in K-562 from CCLE. Cell type proportions are weighted by respective bulk RNA-seq technology detection sensitivity to allow total RNA and mRNA single-cell protocols to be comparable. Single cells were downsampled to 150k reads/cell. **i)** Circos plot showing genomic alterations of known gene fusions (CCLE) in K-562 recovered by STORM-seq.

A limitation of current scRNA-seq methods is data sparsity, making it difficult to differentiate between technical dropouts or true absence of signal at baseline expression levels. STORM-seq’s high sensitivity enables confident interrogation of the heterogeneity of endogeneous TE and TE-derived transcripts in single cells. We examined TE-derived transcript candidates previously discovered with polyA-based bulk RNA-seq in K-562^21^. STORM-seq replicated 76% (25/33) of these candidate TE-derived transcripts. Notably, only four TE-derived transcripts (12%) were expressed in nearly all cells, suggesting that the majority of TE-derived transcripts are heterogeneously expressed even in homogeneous cell populations (**Fig 2c**). Next, we identified additional candidate TE- derived transcripts in K-562 that display cellular heterogeneity (**Methods**), likely missed in previous bulk RNA-seq analyses due to the use of oligo(dT) approaches. STORM-seq detected 40 additional TE-derived transcripts that are also found in bulk total RNA-seq, many of which are associated with lincRNAs and oncogenic processes (**Fig 2c**). As an example, STORM-seq identifies *LTR1A2-PURPL* (P53 Upregulated Regulator Of P53 Levels) as a putative TE-derived long non-coding RNA that spans nearly 300kb, with overlap of similarly long annotated *PURPL* transcript isoforms. *PURPL* lncRNA expression appears to be predominately driven by the TE-derived transcript isoform variants in K-562, also demonstrating the ability of STORM-seq as a tool to dissect isoform level differences in single cells (**Fig 2d**). With *PURPL* being a proposed regulator of p53, we observe the expected anticorrelation of *PURPL* expression and p53 RNA expression with the majority of cells expressing either gene (**Supp.** Fig. 6b).

We expected STORM-seq to be able to capture non-polyadenylated eRNA transcripts in distal enhancers in K-562 single cells. Coverage and detection rates were computed across annotated distal enhancers by GRO/PRO-cap in K-562 cells, as previously described (N=33,207, **Methods**)^28^. Indeed, STORM-seq captures bidirectional expression profiles for detected eRNAs, resembling those from PRO-cap^47^ and TT-seq^26^ (**Fig. 2f**). Compared to VASA-seq, STORM-seq has an ∼2x greater median detection rate of eRNAs in GRO/PRO-cap annotated distal K-562 enhancers^28^ per cell (STORM-seq: 202 enhancers/cell; VASA-seq: 104 enhancers/cell; **Fig. 2g**). Further, we reasoned that given K-562 is a relatively homogeneous cell population, on average, similar enhancers would be utilized and expressed as eRNAs between cells. STORM-seq recovers more shared enhancer/eRNA expression across all bins compared to VASA-seq (**Supp.** Fig. 7a). However, shared enhancer usage did not describe the full complement of detected eRNAs across single cells. As an example, multiple enhancers found within an enhancer cluster upstream of *FTH1* (Ferritin Heavy Chain 1), display differential upstream enhancer usage of a gene involved in iron homeostasis and other oncogenic processes in K-562 (**Supp.** Fig. 7b)^48^. Taken together, STORM-seq enables the ability to dissect eRNA expression profiles in single cells.

One powerful aspect of RNA-seq is the ability to detect fusion transcripts. Some fusions (e.g. *BCR-ABL1* in chronic myeloid leukemia) are pathognomonic for disease^49^. The Cancer Cell Line Encyclopedia (CCLE) has made a catalog of known gene fusions using bulk RNA-seq across many cancer cell lines, including K-562^50^. Using the CCLE K-562 known fusion set, we benchmarked the detection rates of expected gene fusions across bulk RNA-seq, STORM-seq, VASA-seq, and SS3x. We subsampled to 150k reads/cell across scRNA-seq technologies and analyzed the data using STAR-Fusion^51^, as it was the same fusion detection approach in the CCLE. Directly comparing polyA-based methods and total RNA-seq methods for fusion detection rates is complicated due to technical differences in library construction. We reasoned that using full-depth bulk RNA-seq observed fusion fragments per million (FFPM) read support within the same fusion would reduce library construction strategy differences in fusion detection rates (**Methods**). For example, the *BAG6-SLC44A4* gene fusion is expected to have ∼6x the read support in polyA-based RNA-seq than total RNA-seq, while *BCR-ABL1* is approximately equal across library construction strategies. Across scRNA-seq technologies tested, STORM-seq showed the highest proportion of cells containing known gene fusions in K-562, followed closely by SS3x (**Fig. 2h, Supp.** Fig. 7c). VASA-seq consistently had the lowest overall detection rates with *BAG6-SLC44A4*, *C16orf87- ORC6*, and *IMMP2L-DOCK4* not detected (**Fig. 2i, Supp.** Fig. 7d).

### Application to primary human fallopian tube epithelium

We applied STORM-seq to primary human benign distal, pre-menopausal fallopian tube epithelium (FTE) (**Fig. 3**). This tissue is thought to harbor the cell of origin for most high-grade serous ovarian carcinomas (HGSOC)^52^. Therefore, characterization of the repertoire of cell types, states, and drivers of lineage fates found within the benign fallopian tube is critical for our understanding of molecular events leading to HGSOC oncogenesis. Recent scRNA-seq work has proposed different potential progenitor populations and divergent differentiation models^32,33,53,54^. Additionally, intermediate/transitioning cells connecting proposed progenitors to differentiated cell types are largely absent in current studies. To investigate whether STORM-seq can better reconstruct the normal FTE developmental trajectory and identify intermediate/transitioning cell types, we profiled primary pre-menopausal FTE from two donors. We recovered the expected differentiated non-ciliated secretory epithelial cells (NCSE) and ciliated epithelial (CE) cells based on established marker gene sets^33,54^ (**Supp.** Fig. 8). Further, STORM-seq reveals intermediate/transitioning cells along continuous differentiation trajectories to NCSE and CE cell types, and recovers more features per cell, compared to prior studies (**Supp.** Fig. 8-10). Interestingly, within the STORM-seq data, we found a cluster of “dual-feature” cells across donors that simultaneously expressed endothelial (*PECAM1*/*CD31*) and epithelial (*EpCAM*) marker genes (**Supp.** Fig. 11). Given that STORM-seq leverages index sorting, we confirmed that these cells expressed EpCAM^+^ on the cell surface, and were unlikely to be doublets of epithelial and endothelial cells, with *EpCAM* gene expression lower relative to both NCSE and CE populations (**Supp.** Fig. 11-12). We then identified marker genes across cell types, and found progenitor-like gene programs were enriched in these “dual-feature” cells (**Supp.** Fig. 9). Next, we inferred cell trajectories using RNA velocity^55,56^, latent time^56^, pseudotime^57,58^, and principal curves^59^ (**Fig. 3a-e, Supp.** Fig. 13a-c). Notably, the trajectories were consistent across methods and donors, showing a bifurcating lineage from the “dual-feature” cells, along intermediate/transitioning cell types, to NCSE and CE populations (**Fig. 3a-e**, **Supp.** Fig. 13a-c). Therefore, we propose to call this population of cells as “unclassified fallopian tube progenitors” (UCFP). This bifurcating trajectory gives evidence against the de-differentiation model passing through a *RUNX3* or *CD44* progenitor, and supports a common progenitor population that may directly give rise to NCSE and CE cells, based on our analyses (**Fig. 3**, **Supp.** Fig. 13)^32^. Further, we identified potential driver genes shaping lineage fate and identified several transcription factors that are consistent with known FTE biology. Specifically, *PAX8* was found for NCSE lineage commitment – a known marker gene for NCSE cells^33,54^. Within the CE cell lineage, we identified *ULK4*, which has been shown to be important for ciliogenesis^60^ (**Supp.** Fig. 13d-e). As an additional example, two long-intergenic non-coding RNAs (*LINC0188* and *LINC00937*) were inferred as putative drivers for NCSE and CE lineage fates (**Supp.** Fig. 13d-e). To assess the influence of transposable elements (TEs) on lineage commitment in primary human fallopian tube (FT), we examined locus-level expression profiles of LINEs, SINEs, and LTRs, and observe lineage-restricted TE expression along inferred differentiation trajectories from UCFP to ciliated cells across donors (**Fig. 3f**, **Supp.** Fig. 13g). As examples, two LINE elements (L1PA3 and L1ME1) are shown to have expression restricted to the ciliated lineage, and are found within ∼1kb of genes known to participate in cilia beating and ciliogenesis (*IQUB* and *FAM216B*).

**Figure 3.**
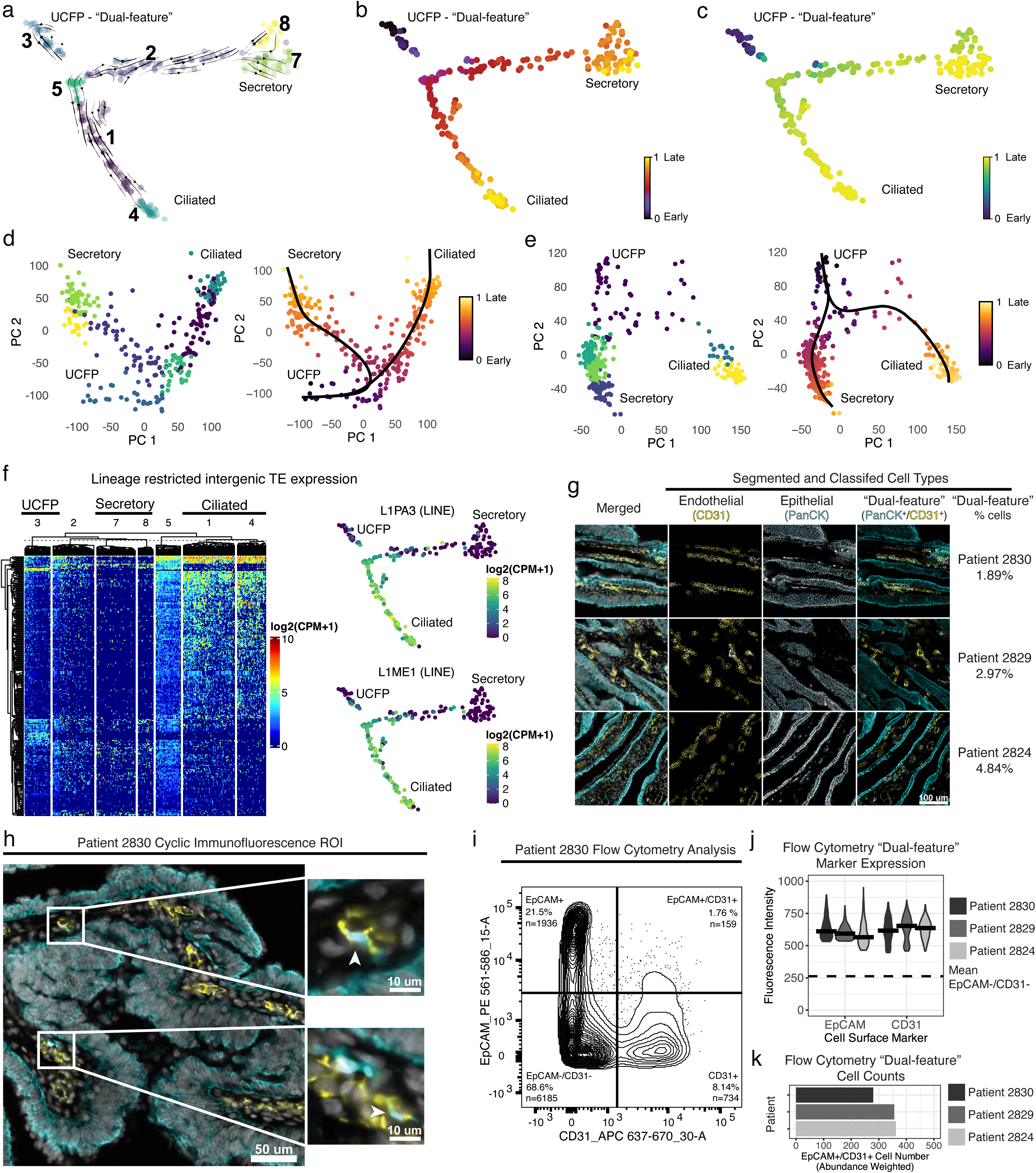
STORM-seq identifies a continuous differentiation trajectory from an unclassified progenitor/”dual-feature” cell population in primary human fallopian tube epithelium. **a)** RNA velocity supports a continuous differentiation process from an unclassified fallopian tube progenitor (UCFP) cells (cluster 3) to an intermediate branch point (cluster 5) to terminally differentiated secretory and ciliated cells (clusters 7-8 and 4, respectively). Thickness and direction of arrows indicate velocity. **b)** Latent time inference of cellular differentiation and **c)** CytoTRACE pseudotime supports a similar differentiation trajectory from early UCFP/”dual-feature” cells towards late, secretory and ciliated cell types. **d)** PCA space embedding of patient 1 with fitted principal curves and pseudotime show similar lineage trajectories as shown above. **e)** PCA space embedding of patient 2 with fitted principal curves and pseudotime show consistency with patient 1 lineage fate from UCFP to differentiatied secretory and ciliated cell types. RNA velocity, latent time, CytoTRACE pseudotime were calculated using scvelo. densMAP embeddings shown for patient 1 (a-c). Principal curves and pseudotime for patients 1 and 2 were calculated using slingshot (d-e). **f)** Representative cell-type specific intergenic transposable element (TE) expression from patient 1 demonstrates lineage restricted TE expression, with example locus-level LINE expression within the ciliated cell lineage. **g)** Independent validation of the presence of the “dual-feature” UCFP cells using cyclic immunofluorescence across 3 additional patients. Cell type classification and quantification of “dual-feature” cells as a proprotion of total cells. **h)** Separate regions of interest (ROI) with zoomed insets to show representative simultaneous expression of epithelial (PanCK) and non-epithelial (CD31) cells. **i-k)** FACS analysis of matched patient single cell suspensions using the same antibodies used for STORM-seq patients 1 and 2 demonstrate similar proportions of UCFP/”dual-feature” cell populations (EpCAM+/CD31+).

Taken together, these results highlight the advantage of high-resolution total RNA profiling for cell type/state and lineage inference. To validate the presence of UCFP cells within the primary human fallopian tube (FT), we subjected full-thickness (epithelial and non-epithelial cells) tissue sections and matched single cell suspensions from three additional donors to cyclic immunofluorescence (CycIF) and flow cytometric analysis. Tissue sections and cell suspensions were stained with an antibody cocktail to delineate epithelial (PanCK or EpCAM) and endothelial (PECAM1/CD31) cell types (**Methods**). Indeed, across all three donors, we observe the presence of epithelial, endothelial, and importantly, the UCFP “dual-feature” cell types at similar frequencies observed in the STORM-seq data (**Fig 3g-k, Supp.** Fig. 14). Thus, by applying STORM-seq to primary human FT tissue, we demonstrate the utility of this method to illuminate new biology and the importance of total RNA profiling in this tissue.

## Discussion

Over the last decade, many scRNA-seq methods have been developed, including both plate-based and droplet protocols, with the primary goal of capturing the whole transcriptome of each cell^61^. Depending on the method being presented, either droplet or plate-based alternatives are cast in a negative light, typically highlighting shortcomings in throughput, gene count, and cost. We believe this is a false dichotomy and choosing the right tool for the job is important depending on the experimental question at hand. While droplet-based protocols are valuable for cataloging cell types in tens to hundreds of thousands of cells, plate-based protocols like STORM-seq, excel at capturing high-resolution, full gene body length profiles of single-cell transcriptomes in hundreds to thousands of cells, where cell states are of primary interest. Choosing a method that is best for a project requires a balance of experimental goals, time, and cost considerations – STORM-seq was developed with these criteria in mind.

The importance of timing in a scRNA-seq protocol has largely been overlooked. The ability to quickly move from dissociated tissue/single cells, to cDNA, to prepared libraries, sequencing, and analysis is paramount for data quality and sample throughput^62^. STORM-seq was designed to minimize manipulations to the cell and its content prior to reverse transcription (RT). Once single cells have been sorted into a plate, only 5 minutes of time is spent before cells undergo reverse transcription. This includes a 3-minute cell lysis/RNA fragmentation step and a 2-minute cooling step. Although the Fragmentation Buffer contains RNase inhibitor, rapidly moving from cells to cDNA is crucial for capturing short-lived RNA transcripts (i.e., eRNAs), as well as limiting RNA degradation^63^. Minimizing cell lysis time ensures that the nuclear membrane remains intact, preventing genomic DNA (gDNA) contamination within libraries, which is particularly detrimental for random hexamer primed protocols like STORM-seq^10^. Indeed, STORM-seq has comparable background genomic alignment rates to bulk total RNA-seq (random hexamer primed) and Smart-seq3xpress (oligo(dT) primed), and in sharp contrast to VASA-seq and Smart-seq-total. Minimizing genomic background alignments is critical as non-RNA species present in the library undermines unspliced transcript abundance estimates, as well as gene biotype detection diversity^64,65^. STORM-seq represents the fastest single-cell total RNA-seq protocol to date, going from single cells to sequence-ready libraries in a single working day.

All sequencing protocols have bias, which can be limited with careful methodological choices during library preparation. STORM-seq rapidly moves from single-cells to a barcoded pool, avoiding cleanup steps before pooling mimicking a bulk RNA-seq approach. Current scRNA-seq methods use bead-based cleanups, including STORM-seq. At every bead-based cleanup step, it is expected to lose a large proportion of the input material, typically requiring additional rounds of PCR amplification to increase yield and potentially introducing biased transcript representation. Therefore, STORM-seq includes unique molecular identifiers (UMIs) to overcome PCR amplification bias (**Fig. 1a-b**). The methodological implementation of introducing the UMI sequences themselves can result in additional technical bias^37^. Prior work has demonstrated that careful design of the UMI-TSO is critical to mitigate strand invasion artifacts^5,8^. Strand invasion occurs when a complimentary strand binds the template switching oligo, causing the subsequent double stranded cDNA to have originated from two different RNA molecules creating a chimeric transcript^13^. STORM-seq has low strand invasion, similar to Smart-seq3xpress and reduced UMI bias, in contrast to competing methods.

STORM-seq is a random hexamer priming protocol, profiling both polyadenylated and non-polyadenylated RNA transcripts (e.g. total RNA), whereas oligo(dT) priming methods measure polyadenylated transcripts either by preferential amplification (Smart- seq3xpress) or introduction of polyA tails to capture total RNA (e.g. VASA-seq and Smart-seq-total). Profiling total RNA expands our insight into unspliced transcripts and non-polyadenylated regulatory features within single-cells, such as transposable elements (TEs), eRNAs, and other transcribed regulatory elements, which are known to play critical roles in cell state and fate. An additional benefit to random hexamer priming is that it is robust to degraded RNA, capturing transcripts that may be lost with oligo(dT) priming methods^41^. To the best of our knowledge, another unique design innovation for STORM-seq is introduction of spike-in ERCC transcripts at the RT and addition of UMIs step, instead of at the cell lysis and RNA fragmentation steps. By taking this approach, it minimizes the skewing of expected ERCC copy numbers^66^. ERCC and related molecular spike-ins provide a means to normalize total RNA content differences, enabling the ability to account for both technical and biological cellular RNA content differences, such as those that can arise during cell replication, *MYC* amplification, and cellular differentiation^15^. ERCC spike-in transcripts are added at known copy numbers, enabling an estimate of assay sensitivity for single or near-single molecule detection. Indeed, STORM-seq accurately reconstructs expected ERCC copy numbers and sensitively detects single copy spike-ins. With this level of sensitivity, STORM-seq can profile total RNA, including short-lived, low expression transcripts (e.g. eRNAs) more robustly than VASA-seq and Smart-seq-total.

RNA velocity^55,56^ and cellular trajectory inference^57,58^ are common analyses in scRNA-seq. RNA velocity infers gene expression dynamics, and ultimately cellular trajectory, through spliced and unspliced transcript abundances, and has been previously shown that single-cell total RNA-seq methods improve RNA velocity inference^6^. We applied STORM-seq to the pre-menopausal human fallopian tube epithelium (FTE), as the stem/progenitor population has remained elusive and is of great interest for reproductive and cancer. Prior efforts utilizing 10x Genomics scRNA-seq proposed divergent models^32,33^. Our results and another prior study both showed that evidence is lacking for the ‘dedifferentiation’ model, and that *RUNX3* does not mark an intermediate population, but is instead, likely an immune cell population^33^. This may be due in part to technological limitations, compounded with the inclusion of confounding cell types (*RUNX3* expressing immune cells) as part of the pseudotime inference^32,59^. Moreover, we observe that *CD44* alone is not a sufficient marker for progenitor-like cell populations. Recent work has proposed that both stroma and ciliated cells both come from mesenchymal/epithelial ‘dual-feature’ cells^33^, similar to those discovered in our study – referred to as unclassified fallopian tube progenitor-like (UCFP) cells, here. In this model, secretory cells had a different, unidentified origin outside of the ‘dual-feature’ cells. Our results show that the two types of epithelial cells likely derive from the same progenitor cell population. Notably, lineage reconstruction is consistent across donors, read depths, and cell lineage fate inference methods. We provide evidence for the first time that these ‘dual-feature’ UCFP populations are also found in whole tissue sections of full thickness human FT. Both secretory and ciliated populations have substantial heterogeneity, particularly within the secretory, non-ciliated epithelial cell lineage. What is traditionally treated as one cell type (secretory cells) likely contains multiple populations of non-ciliated epithelial cells^32,33,53,54^. We also demonstrate the utility of STORM-seq to dissect the contribution of non-coding transcript expression, previously uncharacterized in human FT through the observation of lineage restricted TE expression in ciliated cells. These TEs possess zinc finger protein binding sites, including KRAB-zinc finger and C2H2 zinc finger sites. KRAB-zinc finger proteins are recognized to play important roles in regulating TE expression and demonstrates the power of STORM-seq to simultaneously profile TE and KRAB-zinc finger expression in single cells^67^. Taken together, we show evidence for a putative common progenitor-like cell type (UCFP) that may give rise to both epithelial and endothelial compartments within the human FT.

Finally, while much attention has been given to calculating the cost per cell for scRNA-seq methods, numerous additional factors drive the cost of a sequencing project, rendering this metric incomplete at best^68^. For example, while investment in the liquid handlers required for other methods can cost tens to hundreds of thousands of dollars, this value is not often calculated as part of the price per cell. While STORM-seq is amenable to the use of automation, a simple multi-channel pipette is all that is required for library generation. Additionally, the price per cell is calculated using the number of cells that began the library generation protocol, not the number of cells that are usable after sequencing. STORM-seq recovers nearly all cells after sequencing, limiting the amount of money wasted by generating and sequencing failed libraries. Moreover, STORM-seq is accessible as a commercially available kit, reducing the need to purchase and generate additional reagents for library preparation. Computationally, STORM-seq has been added as a preset within the kallisto|bustools^42^ suite and usable with STARsolo^43^ through synthbar (**Methods**), further reducing the need for custom processing scripts, in the hope of lowering the barrier to entry for this method.

In conclusion, STORM-seq is a random hexamer primed, ribo-reduced RNA-sequencing protocol that does not require specialized equipment, producing the highest complexity single-cell libraries to date. Further, STORM-seq is currently the only single-cell method that can accurately reconstruct TE expression profiles similar to bulk total RNA-seq. It can capture transcribed regulatory elements like eRNAs, and clinically relevant gene fusions in single cells. Finally, when STORM-seq was applied to the human FT, we identified a putative progenitor population expressing both epithelial and endothelial markers, and orthogonally validated in whole tissue section cyclic immunofluorscence (CycIF), combined with flow cytometry. With its carefully optimized design, along with its efficiency and sensitivity in profiling total RNA, we believe STORM-seq represents the state of the art in single-cell total RNA sequencing.

## Materials and Methods in separate document Data and Code availability

STORM-seq K-562, HEK293T, and RMG-2 raw data, SS3x K-562 raw data, and STORM-seq fallopian tube counts have been deposited as GSE181544. Raw FASTQ files for the human fallopian tube epithelium data will be deposited in dbGaP as controlled access at time of publication. Flow cytometry data will be deposited in FlowRepository and will be made available as above. All code and related R and python objects will be deposited in zenodo.

## Supporting information

Supplemental Table 1: Methods timing

Supplemental Table 2: Donor metadata

Supplemental Table 3: TE transcript quantification

## Acknowledgements

We would like to acknowledge all members of the Shen and Triche labs as well as the Van Andel Institute Flow Cytometry, Genomics, and Optical Imaging Core facilities. We would like to thank Takara Bio USA for providing resources for protocol development. This research is supported by National Cancer Institute grant R37CA230748 to HS, Ovarian Cancer Research Alliance fellowship 891749 to BKJ, philanthropic grants from the Grand Rapids Community Foundation, Folz Family Foundation, and Michelle Marie Lunn Hope Foundation to TJT, and startup funds from the Van Andel Institute to TJT and HS. We would like to thank Motion City Soundtrack for the soundtrack to the development of STORM-seq.

