## Supplementary material for "Efficient profiling of total RNA in single cells with STORM-seq": Materials and Methods

### Cell lines and culture conditions

The K-562 cell line was purchased from ATCC (CCL-243). Cells were cultured in RPMI 1640 medium supplemented with 10% FBS and 1% Penicillin/Streptomycin (Gibco). HEK293T cells were purchased from the American Type Culture Collection (ATCC; CRL-3216) and cultured in DMEM medium supplemented with 10% FBS, 2 mM L-glutamine, and 1% Penicillin/Streptomycin (Gibco). RMG-2 cells were purchased from the JCRB Cell Bank and Seikiusi XenoTech, LLC. RMG-2 cells were maintained in DMEM/F12, supplemented with 10% FBS and sub-cultured at 1:3-1:5 every 7-9 days. Fresh media was added every 2-3 days between sub-culturing.

### Human Fallopian tube epithelium sample acquisition

Fallopian tube fimbriae were collected at Spectrum Health Butterworth Hospital (now Corewell Health) in Grand Rapids, Michigan. After initial screening for benign conditions not affecting the epithelium of the fallopian tube (e.g. uterine fibroids), patients undergoing total hysterectomy with salpingo-oophorectomy were provided written informed consent for sample collection and analysis. The study was approved by Institutional Review Board (IRB) of the Van Andel Research Institute under IRB #19017. Participant enrollment and specimen collection was conducted via partnership with the Corewell Health biobank program, SHARE Biorepository, approved the Corewell Health West IRB (#2017-198). Following surgical resection, the tissue was placed in saline and the single-cell dissociation protocol was started within 20 minutes of receiving the tissue. Patient metadata can be found in **Supplemental Table 2**.

### Fallopian tube epithelium dissociation to single cells

Dissociation of fresh Fallopian tube epithelium was done using the Milltenyi gentleMACS Octo Dissociator with heaters and the human Tumor Dissociation Kit (Miltenyi Biotec). Briefly, the tissue was manually minced in a sterile Petri dish. Minced tissue was placed into Milltenyi gentleMACs C tubes, complete with 4.7 mL of DMEM (Gibco) and supplemented with enzymes A, H, and R at the manufacturer recommended working concentrations (Miltenyi Biotec). Next, the tissue was dissociated into a single-cell suspension using the “soft tumor type” program (37C_h_TDK_1) on the gentleMACS Octo Dissociator. Following conclusion of the dissociation program, sample tubes were centrifuged at 300 x g for 3-5 minutes to pellet the cells. The supernatant was removed and the pellet resuspended in filter-sterilized flow sorting buffer (HBSS + 2% FBS + 25 mM HEPES). Remaining intact tissue was removed by passing the suspension through a Milltenyi SmartStrainer (Miltenyi Biotec) into a new 50 mL Falcon tube (Corning). Viable cells were counted using Trypan blue and the BioRad TC20 or Thermo Fisher Countess 3 Cell Counter. The remaining, strained single-cell suspension was transferred to 2 separate 15 mL Falcon tubes (Corning) and centrifuged to pellet cells (300 x g for 3-5 minutes). Pellets were resuspended in freezing media (90% FBS + 10% DMSO) and cryopreserved.

**Cell line flow cytometry sorting for STORM-seq**

Cell lines were spun down at 300 x g for 5 minutes at 4C and washed in Flow Buffer (HBSS with no divalent cations + 2% FBS + 25 mM HEPES). Prior to sorting, cell line suspensions were resuspended in Flow Buffer with 0.5 ug/mL DAPI for active viability surveillance during sorting. Single, live cells were index sorted into each well of a 384-well plate containing Fragmentation Buffer (Takara) using a BD FACSymphony S6 sorter running BD FACSDiva v9.1.4 and equipped with BD StepSort. The S6 was run with a 130 um nozzle at 14 psi. For cell deposition into 384-well plates or bulk, we used single cell sort mode and purity, respectively. Representative, full gating schemes for K-562, HEK293T, and RMG-2 can be found in **Supp. Figs. 15-17**.

**Fallopian tube cell sorting and analysis**

Cryopreserved primary cells were rapidly thawed in a 37C water bath. The thawed cells were gently resuspended in several volumes of Flow Buffer (HBSS with no divalent cations + 2% FBS + 25 mM HEPES) and pelleted at 300 x g at 4C for 5 minutes. Cell pellets were resuspended in 1 mL of Flow Buffer and viable cells were counted using trypan blue and either the BioRad TC20 Automated Cell Counter (BioRad) or Countess 3 Cell Counter (Thermo Fisher). Cells were pelleted and resuspended in 100 uL of Flow Buffer and stained for 60 minutes on ice, in the dark with EpCAM-PE (CD326; clone 1B7; Invitrogen; 1:100), CD235a-FITC (clone HIR2 (GA-R2); Invitrogen; 1:100), and CD31-APC (clone WM-59/WM59; Invitrogen; 1:100). Cells were pelleted and washed twice with 1 mL flow sorting buffer, with the final wash resuspending cells in 300-500 uL of Flow Buffer. To assess viability, DAPI (Thermo Fisher) was added to a final concentration of 2 μg/mL (Beckman Coulter MoFlo Astrios) or 0.5 ug/mL (BD FACSymphony S6) to stained cells without washing for active viability surveillance throughout sorting. For STORM-seq of primary fallopian tube cells, single, live, EpCAM$^{+}$, CD235a$^{-}$ cells were index sorted using a Beckman Coulter MoFlo Astrios cell sorter running Summit v6.3, into designated wells of a 384-well plate containing Fragmentation Buffer (Takara). The Astrios was equipped with a 100 um nozzle at 25 psi. Abort mode was set to single and drop envelope to 0.5. For analysis of additional fallopian tube samples, single cell suspensions were processed as described above and analyzed using a BD FACSymphony S6 cell sorter. Details of the S6 are the same as described above but running BD FACSDiva v9.5.1. The antibodies plus fluorochromes and viability dyes for cell sorting used the following excitation/emission wavelengths and associated filter sets: DAPI 355-450/50, CD235a-FITC 488-515/50, EpCAM-PE 561-586/15, and CD31-APC 637-670/30. A representative, full gating strategy can be found in **Supp. Fig. 18**.

**Bulk total RNA isolation and library construction from K-562 cells**

Total RNA from 5 million K-562 cells were isolated using TRIzol (Invitrogen). Libraries were prepared by the Van Andel Genomics Core from 500 ng of total RNA using the KAPA RNA HyperPrep Kit with RiboseErase (v1.16) (Kapa Biosystems, Wilmington, MA USA). RNA was sheared to 300-400 bp. Prior to PCR amplification, cDNA fragments were ligated to Illumina UDIs. Quality and quantity of the finished libraries were assessed using a combination of Agilent DNA High Sensitivity chip (Agilent Technologies, Inc.) and QuantiFluor® dsDNA System (Promega Corp., Madison, WI, USA).

**STORM-seq library construction for cell lines**

Single cells were prepared by hand pipetting using the STORM-seq kit and protocol (Takara Bio Cat Number 634751, detailed protocol: https://doi.org/10.5281/zenodo.15178455). To begin, cells were sorted into a 384-well plate (Eppendorf twin.tec PCR Plate 384 LoBind) containing 2.17ul Fragmentation Buffer (1.17ul PBS pH 7.2 [Gibco 20012-027], 0.17ul 10X Lysis Mix, 0.17ul SMART scN6, 0.69ul scRT buffer) and immediately transferred to a thermocycler, heated for 3 minutes at 85C, and cooled on ice for 2 minutes. Once cooled, 1.17ul First Strand Master Mix (0.75ul SMART scTSO mix, 0.08ul RNase Inhibitor, 0.34ul SMARTscribe RT) is added to each well. Immediately prior to dispensing, ERCC RNA Mix 1 is added to the First Strand Master Mix to create a 1:1,000,000 dilution. First Strand Synthesis is performed at 42C for 180 minutes, followed by 70C for 10min and 4C hold. After 1st Strand Synthesis, PCR1 (10 cycles of amplification: 94C 1 min, [98C 15s, 55C 15s, 68C 30s]x10, 68C 2 min, 4C hold) was performed with the addition of SMARTer RNA Unique Dual Index Sets A-D (SMARTer® RNA Unique Dual Index Kits 96U sets A-D, Takara Bio Cat Numbers 634752, 634753, 634754, 634755). 4.67ul of PCR1 Master Mix (0.33ul nuclease free water, 4.16ul SeqAmp CB PCR buffer, SeqAmp DNA Polymerase) is added to each well. Each well then receives 1ul of a Unique Dual Index. UDIs were diluted 1:4 in 10mM Tris-HCl, pH 8.0 (Teknova) prior to addition. The plate was pooled and a bead-based cleanup (Beckman Coulter AMPure XP Beads) was performed. 162ul of rRNA Depletion Master Mix (123.12ul nuclease free water, 16.2ul 10X ZapR buffer, 11.02ul scZapR, 11.02ul sc-R Probes) is used to elute the depleted cDNA from the dried beads. The eluate is incubated at 37C for 60 min, 72C for 10 min, 4C hold. Finally, 12 cycles of amplification (94C 1 min, [98C 15s, 55C 15s, 68C 30s]x12, 4C hold) were performed for PCR2 (PCR2 Master Mix (208ul nuclease free water, 400ul SeqAmp CB PCR Buffer, 16ul PCR2 primers, 16ul SeqAmp DNA polymerase) and the final library was eluted in 20ul 10mM Tris-HCl pH 8.0. After QC, where necessary, an additional bead clean-up was performed to remove any remaining adapter-dimer.

**STORM-seq library construction for fallopian tube**

Single cells were prepared using the SPT Labtech Mosquito (HV) using the STORM-seq kit and protocol (Takara Bio Cat Number 634751, detailed protocol: https://doi.org/10.5281/zenodo.15178455). To begin, cells were sorted into a 384-well plate (Eppendorf twin.tec PCR Plate 384 LoBind) containing 2.17ul Fragmentation Buffer (1.17ul PBS pH 7.2 [Gibco 20012-027], 0.17ul 10X Lysis Mix, 0.17ul SMART scN6, 0.69ul scRT buffer) and immediately transferred to a thermocycler, heated for 3 minutes at 85C, and cooled on ice for 2 minutes. Once cooled, 1.17ul First Strand Master Mix (0.75ul SMART scTSO mix, 0.08ul RNase Inhibitor, 0.34ul SMARTscribe RT) is added to each well. Immediately prior to dispensing, ERCC RNA Mix 1 is added to the First Strand Master Mix to create a 1:1,000,000 dilution. First Strand Synthesis is performed at 42C for 180 minutes, followed by 70C for 10min and 4C hold. After 1st Strand Synthesis, PCR1 (10 cycles of amplification: 94C 1 min, [98C 15s, 55C 15s, 68C 30s]x10, 68C 2 min, 4C hold) was performed with the addition of SMARTer RNA Unique Dual Index Sets A-B (SMARTer® RNA Unique Dual Index Kits 96U sets A-B, Takara Bio Cat Numbers 634752, 634753). 4.67ul of PCR1 Master Mix (0.33ul nuclease free water, 4.16ul SeqAmp CB PCR buffer, SeqAmp DNA Polymerase) is added to each well. Each well then receives 1ul of a Unique Dual Index. UDIs were diluted 1:3 in 10mM Tris-HCl, pH 8.0 prior to addition. The plate was pooled into 4 tubes of 96 cells, each containing a unique set of UDIs and a bead based cleanup (Beckman Coulter AMPure XP Beads) was performed. 41ul of rRNA Depletion Master Mix (30.78ul nuclease free water, 4.05ul 10X ZapR buffer, 2.76ul scZapR, 2.76ul sc-R Probes) is used to elute the depleted cDNA from the dried beads. The eluate is incubated at 37C for 60 min, 72C for 10 min, 4C hold. Finally, 12 cycles of amplification (94C 1 min, [98C 15s, 55C 15s, 68C 30s]x12, 4C hold) were performed for PCR2 (PCR2 Master Mix (52ul nuclease free water, 100ul SeqAmp CB PCR Buffer, 4ul PCR2 primers, 4ul SeqAmp DNA polymerase) and the final library was eluted in 20ul 10mM Tris-HCl pH 8.0. After QC, where necessary, an additional bead clean-up was performed to remove any remaining adapter-dimer.

**STORM-seq sequencing**

The manually prepared K-562 single cells used to assess fragmentation time and coverage, were pooled and sequenced using 2x75 bp sequencing on an Illumina NextSeq 500 (Illumina Inc., San Diego, CA, USA). Base calling was done by Illumina NextSeq Control Software (NCS) v2.0 and output of NCS was demultiplexed and converted to FASTQ format with Illumina Bcl2fastq v1.9.0. Bulk libraries were pooled and 2x50 bp sequencing was performed on an Illumina NovaSeq 6000 sequencer using a 100 cycle S2 sequencing kit (Illumina Inc., San Diego, CA, USA). The HEK293T, RMG-2, and K-562 mixture was sequenced through the VAI Genomics Core on a NovaSeq 6000 sequencer using a 2x150 cycle sequencing kit, across two lanes. Patient 1 fallopian tube epithelium STORM-seq libraries were also sequenced on a NovaSeq 6000 sequencer using a 2x100 bp sequencing kit. Patient 2 fallopian tube epithelium STORM-seq libraries were sequenced on a Hiseq 4000 using a 2x150 bp sequencing kit at Fulgent Genetics. Base calling was done by Illumina NovaSeq Control Software (NCS) v2.0 and output of NCS was demultiplexed and converted to FASTQ format with Illumina Bcl2fastq v1.9.0. Base calling for the HiSeq sequencing run was performed at Fulgent Genetics and returned as FASTQ files.

**VASA-seq library preparation**

Single, viable passage-matched HEK293T and K-562 cells were sorted as described above into 384-well plates and buffer provided by Single Cell Discoveries (SCD). Plates were then processed as previously described in Salmen et al. 2022, Nature Biotechnology and sequenced using a 1x75 bp kit on a NextSeq 500/550 by SCD. Quality control (QC) metrics provided by SCD support high-quality libraries and sequencing results. In total across both 384-well plates, >92% of single cells had libraries passing QC thresholds with >1000 endogeneous reads, and an average raw read depth of 179,500 reads/cell.

**Smart-seq3xpress library preparation**

Single, viable K-562 cells were sorted into 384-well plates and libraries were generated as previously described in Hagemann-Jensen et al. 2022, Nature Biotechnology. Libraries were sequenced on a NextSeq 2000 using a 300-cycle flow cell. Cells were sequenced to an average read depth of ~380,000 reads/cell.

**Publicly available data sets for benchmarking**

*Cell lines*

The following publicly available datasets were downloaded and used for comparison to STORM-seq: 1) K-562 SMART-seq2 (Illumina runs) - SRP132313, 2) HEK293T SMART-seq-total UMI - GSE151334, 3) HEK293T Smart-seq3xpress - E-MTAB-11467, 4) HEK293T 10x 3’, v3.1 Chromium data were downloaded from the 10x Genomics website, 5) Bulk HEK293T mRNA and total RNA – ERP003460, 6) Bulk K-562 mRNA – SRR8615717, 7) Smart-seq – SRP041736, 8) Bulk RMG-2 mRNA – SRR926261, 9) Bulk K-562 replicates 1 and 2 stranded PRO-cap bigWigs – ENCSR882DWM, 10) Bulk K-562 replicate 1 and 2 stranded TT-seq bigWigs – GSM4610686 and GSM4610687.

*Primary human benign fallopian tube*

The following publicly available datasets were downloaded and used for comparison to STORM-seq: 1) SMART-seq2 – GSE139079, 2) 10x Genomics 3’ – GSE151214

**Demultiplexing Smart-seq3xpress and VASA-seq data**

Unmapped Smart-seq3xpress (SS3x) HEK293T and K-562 BAM files were demultiplexed to individual cell FASTQ files by extracting reads assigned to expected cell barcodes in the CB BAM tag (concatenation of i5 and i7 indices, yielding a 20 bp barcode) and appending conserved sequences to delineate internal and 5’ UMI containing reads. Briefly, to reconstruct the expected 4 FASTQs per cell file structure, cell barcodes (CB BAM tag) were added to index 1 and index 2 FASTQ files, and respective read 1 and read 2 reads to separate FASTQ files. If the UB BAM tag (UMI) was populated, the conserved ‘ATTGCGCAATG’ sequence motif was added, then the UMI (UB tag), followed by the ‘GGG’ motif. Internal and 5’ UMI containing reads from SS3x were separated as previously described, using the conserved ‘ATTGCGCAATG’ sequence motif at the start of read 1 to delineate 5’ UMI reads. VASA-seq data were demultiplexed to single cell FASTQ files using provided scripts from https://github.com/hemberg-lab/VASAseq_2022.

*Trimming single-cell and bulk RNA-seq*
Smart-seq-total HEK293T cells were trimmed using cutadapt (v4.4) following the strategy as previously described. Briefly, read 2 reads were trimmed using default parameters with the following modifications: 1) -j 4, 2) -u 6, 3) -a AAAAAAAAAA, and 4) -m 18. Next, UMI containing reads (read 1) ending in a poly(T) sequence (‘TTT’) were extracted, and matched read 2 FASTQ files were re-paired using seqkit pair (v2.7.0). VASA-seq data were pre-processed and trimmed using cutadapt (v4.4) through provided scripts at https://github.com/hemberg-lab/VASAseq_2022. After trimming, the UMI and cell barcode were appended to the 5’ end of the read. Importantly, in the provided concatenator.py, a conversion error occurs when extracting and appending the UMI and cell barcode where a base quality score of 14 (ASCII character ‘/’) is unintentionally converted to a base quality score of 46 (ASCII character ‘O’). As a result, during the addition of the UMI and cell barcode to the 5’ end of the read, these base qualities were reverted to their original value. SMART-seq K-562 data were trimmed using TrimGalore (v0.6.7) with default parameters. SMART-seq2 K-562 data were trimmed using TrimGalore (v0.6.7). Default parameters were used with the following modifications: 1) --trim-n, 2) --length 36, 3) --paired, and 4) --fastqc. Bulk total RNA-seq of K-562 cells were trimmed using TrimGalore (v0.6.3) using the same set of parameters as SMART-seq2 data.

**Protocol timing**
All protocol timing steps were estimated using published methods sections. Hands on time is defined as actively manipulating the sample with human intervention (e.g., pipetting), and hands off time is where the sample is not actively being handled (e.g., PCR amplification in the thermalcycler). The time spent for each general step (e.g., bead cleanup) was based on Picelli et al., 2014, Nature Protocols. Additionally, Smart-seq3xpress and snapTotal-seq timing analysis was independently verified from authors of their respective method.

**Subsampling bulk and single-cell RNA-seq technologies**

All subsampling was performed using seqtk (v1.4-r122) sample on pre-processed/trimmed reads across technologies, setting the random seed to 100, unless specified otherwise.

**Gene/transcript detection benchmarking**

Pre-processed single-cell and bulk FASTQs were mapped and quantified using Ensembl 101 annotations, supplemented with ERCC or molecular spike-ins where needed.

*STORM-seq*

STORM-seq HEK293T, K-562, and RMG-2 reads were mapped and quantified at the gene and transcript isoform level using kallisto|bustools (kallisto: v0.50.1, bustools: v0.43.2) and kb_python (v0.28.2) with the -x STORM-seq preset. For transcript isoform quantification, kb_python was called with the --tcc option enabled to output transcript compatibility counts (TCC). TCCs were quantified using kallisto quant-tcc to produce transcript isoform level counts. Further, fragment length bias was accounted for in quant-tcc as well as including at least 50 bootstraps for gene/transcript isoform quantification. Per-cell mapping rates were extracted from the log files.

*VASA-seq*

Passage matched (for direct comparison to STORM-seq) K-562 and HEK293T VASA-seq data generated by Single Cell Discoveries were mapped and quantified against Ensembl 101 using kallisto|bustools (kallisto: v0.50.1, bustools: v0.43.2) following the kb_python analysis toolchain, with modifications to support this data type. Briefly, kallisto bus was used with default parameters and 1) -x 0,6,8:0,0,6:0,14,0, 2) --fr-stranded. Next, bus files were sorted (bustools sort), inspected (bustools inspect), and counted (bustools count). For bustools count, default parameters were used with the following modifications, 1) --multimapping, 2) --umi-gene. Next, kallisto quant-tcc was used to quantify transcript abundance with default parameters and 1) -t 8, 2) -l 550, 3) -s 300, 4) -b 50, and 5) --matrix-to-files. The average fragment length and standard deviation parameters were specified based on VASA-seq bioanalyzer traces provided (**Supp. Fig. 19**). Per-cell mapping rates were extracted from the log files.

*Smart-seq-total*

Smart-seq-total HEK293T pre-processed FASTQs were mapped and quantified using a similar strategy as the VASA-seq data, using kallisto|bustools. Briefly, kallisto bus was used with default parameters and the following modifications to accommodate the read structure and library strategy: 1) -x 0,0,7:0,7,23:1,0,0 and 2) --unstranded. To quantify transcript abundances, kallisto quant-tcc was used with the same parameters as VASA-seq, without specifying fragment lengths. Average fragment lengths were not able to be estimated due to lack of available bioanalyzer traces and library read structure. Per-cell mapping rates were extracted from the log files.

*Smart-seq3xpress*

HEK293T and K-562 pre-processed SS3x FASTQs were mapped and quantified using a similar strategy as VASA-seq, using kallisto|bustools. Briefly, per-cell 5' UMI reads and internal reads were extracted as described above, prior to subsampling and mapping/quantification. Next, for gene/transcript quantification, kallisto bus was used with 5' UMI reads with default parameters and the following modifications, 1) -x 0,0,0,1,0,0:2,0,21:2,24,0,3,0,0, 2) --tag ATTGCGCAATG, 3) --paired, 4) --fr-stranded, and 5) -t 8. Transcript abundance quantification was performed using kallisto quant-tcc with default parameters and the following modifications 1) -b 50 and 2) --matrix-to-files.

**Gene body coverage**

In order to compare gene body coverage from total RNA and mRNA protocols, Ensembl 101 protein coding annotations were extracted and converted to BED12 format. Prior to mapping, Smart-seq3xpress 5' UMI and internal reads were separated as described above. Single-cell RNA-seq data containing UMIs were mapped using STARsolo (v2.7.11a) unless noted otherwise. Non-UMI containing data were mapped using STAR (v2.7.11a). Single cell BAMs were merged using samtools (v1.9) merge with default parameters. For the 10x Genomics 3', v3.1 Chromium data, raw FASTQs were downloaded from the 10x Genomics website (10k 1:1 mixture of HEK293T/NIH3T3 3', v3.1 Chromium Controller). Cell Ranger (v7.0.1) was used to align the data to Ensembl 101 with default parameters with the following modification as recommended by 10x for this data: --expect-cells 10000. HEK293T cells were subset from the resulting BAM using the 10x Genomics tool, subset-bam (v1.1.0) through the annotated cell barcodes provided by 10x Genomics with this data set. The subset HEK293T 10x Chromium CellRanger BAM was used directly. The RSeQC (v4.0.0) geneBody_coverage.py script was used to quantify gene body coverage across technologies. Plots were generated using R (v4.4.1) and ggplot2 (v3.5.1).

**Lightweight tool for addition of synthetic cell barcodes**

Synthbar is a lightweight, self-contained program for adding synthetic cell barcodes to sequencing data. By default, it reads an uncompressed or gzip-compressed FASTQ file as input, prepends a cell barcode to each read sequence (with corresponding quality scores), and then writes the modified reads to the computers standard output. As a standalone tool, there are several options to the user, including writing the output straight to a file and modifying the cell barcode prepended to each read. The STORMsolo pipeline, which processes raw STORM-seq data through synthbar and STARsolo, uses the default options for synthbar. The default pipeline will delete the modified FASTQs to reduce disk usage; however, the user can optionally retain these files. https://github.com/jamorrison/synthbar

**Genomic background mapping and poly(A/T) track annotation**

For background genomic alignments, Ensembl 101 gene annotations, repeat masker annotated intergenic TEs, and known R-loops (RLHub Bioconductor/R package, v1.9.0) were combined to establish the "annotated space" reference. All single-cell and bulk total RNA-seq HEK293T samples were downsampled to 100k raw reads per cell using seqtk. Single-cell technologies were mapped using STARsolo (v2.7.11a) and bulk total RNA-seq was mapped using STAR (v2.7.11a) with default parameters and the following modifications. For scRNA-seq samples, relevant UMI, cell barcode, and cDNA fragment structures were used to guide STARsolo parameter specification with: 1) --outSAMtype BAM SortedByCoordinate, 2) --outSAMattributes NH HI nM AS CR UR CB UB GX GN, 3) --soloUMIdedup Exact, 4) --soloMultiMappers EM, 5) --soloFeatures GeneFull, and 6) --soloOutFileNames output/ features.tsv barcodes.tsv matrix.mtx. For bulk total RNA-seq, STAR parameter modifications were: 1) --outSAMattributes NH HI nM AS GX GN and 2) --outSAMtype BAM SortedByCoordinate. Primary alignments were then extracted using samtools (v1.17) and the subcommand view with the option -F 256. BAM files were then name sorted with samtools sort and converted to BED (Smart-seq-total and VASA-seq) or BEDPE (STORM-seq, Smart-seq3xpress, and bulk total RNA-seq) with bedtools (v2.31.0) bamtobed. Next, BED/BEDPE files were intersected using bedtools, with the gene, TE, and R-loop genomic reference, keeping alignments that did not overlap known annotations, merged into a single BED file per-technology, and sorted using bedtools. Merged BED files were then read into R (v4.4.1) using rtracklayer (v1.64.0) and per-cell counts of reads mapping to unannotated space were quantified. Percentages of genomic background mapping was calculated as a fraction of input raw read counts (e.g., 100k reads/cell) and plotted using ggplot2 (v3.5.1). Poly-A/T track generation was performed using a custom Python script based on prior work defining criteria for a genomic poly(A/T) run. Briefly, a greedy algorithm was implemented where the genome was scanned for a minimum of six consecutive nucleotides (e.g. A or T), allowing for up to two consecutive mismatches, and required to have a minimum of 60% A/T content to be considered a poly(A/T) run. Reads mapping to unannotated space were then overlapped based on their genomic coordinates to annotated poly(A/T) runs using GenomicRanges (v1.56.0) and visualized using GViz (v1.48.0).

**UMI bias analysis and simulation of inter-gene collisions**

Single-cell technologies were subsampled to 100k reads/cell in HEK293T using seqtk and mapped to Ensembl 101, including spike-in controls where necessary, using STARsolo as described above. BAM files were filtered, removing secondary and supplementary alignments. UMI sequences were extracted from the UB BAM tag, and merged into a single file with corresponding cell barcodes extracted from the RG BAM tag. UMI bias was calculated as the most abundant (e.g., greatest count) per cell in R (v4.4.1) and plotted using ggseqlogo (v0.2). UMI diversity was calculated by keeping only unique UMI sequences across all cells as a reflection of the original UMI pool diversity in R and plotted using ggseqlogo. UMI frequency across all cells was calculated by keeping all UMIs and plotting the nucleotide diversity at each base pair along the length of the UMI within each technology in R, using ggseqlogo. For UMI inter-gene collision simulations, theoretical UMI space was generated using the expected motifs for each single-cell technology. A number of fixed "genes detected" per cell was used as a proxy for varying read depth mapping/quantification results (e.g., 100, 500, 1000, 2000, 4000, 6000, 8000, 10000, and 12000 detected genes per cell). Empirical values for mean and dispersion parameters were estimated from single-cell technologies using edgeR (v4.4.1) and found to be on average, approximately 6.1 for both mean (6.109) and dispersion (6.173), approximating a Poisson distribution. Thus, all simulations were carried out using empirically derived parameters for both mu and theta, across genes detected, sampling UMIs with replacement. To establish the expected inter-gene UMI collision rate, the number of genes assigned with the same UMI were considered an inter-gene collision. Rates were calculated as the ratio of collision events to the total number of allocated UMIs. Simulation scripts and logs are available at https://github.com/biobenkj/stormseq_protocols. Simulations were run 10 times for each gene detection value, each with a separate random seed (noted in each log file). The observed inter-gene collision rates were generated by subsampling all single-cell technologies to 100k reads/cell and mapping using STARsolo as described above. Given that each technology identifies approximately 8000 genes/cell (**Fig. 1g**), the expected rate from the simulation of 8000 genes was used in calculation of the observed/expected rate.

**Transcript detection robustness**

To assess the ability of each single-cell total RNA-seq technology for transcript isoform detection rate and abundance variability across simulated experiments, we constructed the following sampling framework. First, each single cell across technologies in HEK293T were subsampled with seqtk to the maximum read depth possible to compare all technologies – 150k reads/cell as the upper-bound for generated VASA-seq libraries. Next, 10 random seeds were specified to subsample cells to 50k reads/cell with seqtk as a proxy for 10 separate experiments. These randomly subsampled cells across technologies were then mapped and quantified at the transcript isoform level using kallisto|bustools as described above, including fragment length bias adjustment and 50 bootstraps per cell. Bootstrapped abundance.h5 files were processed in R (v4.4.1) using sleuth (v0.30.1). Within each transcript, subsamples were summarized in R to calculate the detection rate (proportion of bootstrapped TPM > 0), mean transcript expression (weighted by detection rate), and a detection rate-adjusted coefficient of variation. Weighting by detection rate provides a compromise to account for both technical and structural zeroes, enabling meaningful comparisons across cells and subsamples. Summarized metrics across all 10 simulated experiments were then calculated by taking the intersection of commonly detected transcripts across all three technologies, requiring a minimum detection rate of 10% across subsamples. To assess transcript length biases, results were stratified across annotated mRNA transcript lengths (Ensembl 101). Transcripts below 500 bp were excluded due to exclusion of these fragments during bead-based cleanups during library preparation. Plots were generated in R (v4.4.1) and ggplot2 (v3.5.1).

**Sensitivity analysis of single molecules in STORM-seq**

STORM-seq HEK293T, K-562, and RMG-2 were subsampled to specified depths (e.g., 100k, 250k, 500k, and 1M reads/cell) using seqtk. Reads were mapped and quantified using kallisto|bustools as described above, adjusting for fragment lengths and including 50 bootstraps per cell. ERCC spike-ins were extracted across cell types and converted to a SingleCellExperiment (v1.26.0) object in R (v4.4.1). ERCC annotations were downloaded and converted to expected copy numbers based on the 1:1 million dilution. Expected copy numbers of ERCC Spike-in Mix 1 was derived from Pollen et al., 2014, Nature Biotechnology, at 28,000 ERCC molecules per reaction when used at a 1:20,000 dilution.

**Strand invasion quantification**

Strand invasion for STORM-seq and Smart-seq3xpress were calculated as previously described in Hagemann-Jensen et al., 2022 Nature Biotechnology with minor modifications. Briefly, pyTSOfilter (https://github.com/cziegenhain/pyTSOfilter) was expanded to allow it to be used with reverse stranded protocols, like STORM-seq, and specification of the UMI containing read depending on the library architecture (e.g., either read 1 or read 2). Modified scripts can be found at https://github.com/biobenkj/stormseq_protocols. Strand invasion percentages were plotted across cell types and technologies in R with ggplot2.

**Enhancer RNA (eRNA) analysis**

Enhancer RNA analyses were carried out using K-562 nascent RNA GRO-cap/PRO-cap annotated distal transcribed regulatory elements (TREs) from PINTS (https://pints.yulab.org/). TREs greater than 1kb were filtered out (n=1541; min size = 101 bp, mean = 494.4 bp, median = 401 bp, and max size = 3584 bp) and remaining elements were resized to 1kb from their respective centers using GenomicRanges in R to allow for consistent ranges between per cell quantification/detection and profile plots. The distal TREs were then intersected with gene bodies (Ensembl 101) using bedtools intersect, keeping the non-overlapping set of elements (n=9378), and was converted to a GTF for STAR index generation and quantification. STORM-seq and VASA-seq were subsampled to 150k reads/cell using seqtk and mapped using STARsolo as described above, including the distal TREs. For eRNA profile plots, single-cell BAMs were merged using samtools, and split into forward and reverse strand bigWigs using deeptools (v3.5.1) bamCoverage, setting the binSize to 10 bp and normalized using bins per million mapped reads (BPM). Downloaded PRO-cap and TT-seq replicate bigWigs were combined into per-technology bigWigs using deeptools bigWigAverage, setting --binSize 10 for consistency with single-cell data. Next, deeptools computeMatrix reference-point was used to calculate the normalized coverage across distal TREs, adding --referencePoint = center, --outFileNameMatrix , --sortRegions no, --missingDataAsZero, -a 500, -b 500, and --binSize 10. Data matrices were then read into R and plotted with ggplot2. Given that the PRO-cap and TT-seq data were processed and normalized differently, the normalized forward and reverse strand coverages were scaled to be on similar axes. STORM-seq and VASA-seq normalized coverages were also scaled using the same scaling factor of 3. Plots were generated using ggplot2. The number of detected enhancers per cell (UMI count > 0) were calculated using STARsolo as described above. For both STORM-seq and VASA-seq, 100 randomly sampled cells using the sample() function in R were used to compare enhancer detection rates. Cells were binned into 2-5, 5-10, 10-15, and >15 cells sharing common detected enhancers and plotted using ggplot2. Per-cell detection rates were also plotted using ggplot2.

**Gene fusion calling**

K-562 STORM-seq, VASA-seq, bulk total RNA, and Smart-seq3xpress (internal reads) were aligned using the STAR-Fusion pipeline (v1.13.0) and pre-built STAR-Fusion CTAT reference genome GRCh38_gencode_v37_CTAT_lib_Mar012021.plug-n-play with default parameters and the following modifications: --max_sensitivity. Specifically, for SMART-seq-style sequencing runs, recommended best practices were followed according to [https://github.com/STAR-Fusion/STAR-Fusion/wiki/STAR-Fusion-scRNA-seq](https://github.com/STAR-Fusion/STAR-Fusion/wiki/STAR-Fusion-scRNA-seq). Following alignment and joint fusion calling, single-cell fusion calls were deconvolved using the aggregate_and_deconvolve_fusion_outputs.py script provided with STAR-Fusion. Each single cell was then analyzed for fusions. Bulk total and mRNA K-562 were also analyzed using STAR-Fusion default parameters. Fusion Fragments Per Million total reads (FFPM) were calculated from the mRNA and total RNA-seq STAR-Fusion runs within known K-562 gene fusions from CCLE (https://depmap.org/portal/cell_line/ACH-000551?tab=fusions). The ratio of mRNA FFPM and total RNA-seq FFPM were then generated to weight detection rates in mRNA and total sc-RNA-seq fusion results. Ratios were as follows: BAG6--SLC44A4 = 5.99, BCR--ABL1 = 1.02, C16orf87--ORC6 = 2.85, IMMP2L--DOCK4 = 1.31, NUP214--XKR3 = 2.61, UPF3A--CDC16 = 3.00, and XACT--

LRCH2 = 0.90. Proportion plots were plotted with ggplot2 and circos plots were plotted with circlize (v0.4.16) in R.

**TE expression quantification**

Single-cell TE sub-family-level enrichment was calculated as previously described in Shao and Wang, 2020 Genome Research. Briefly, where needed, adapter sequences were trimmed using Trim Galore (v0.6.5). The adapter trimmed reads were then mapped to the reference genome hg38 (Ensembl 101) using STAR or STARsolo as described above, increasing --outFilterMultimapNmax to 500. The mapped reads were assigned to their respective features using featureCounts (v2.0.1) if there was a minimum fractional overlap of 0.1. Multimapped reads were also counted (-M option) but fractional counts were assigned to their respective features (--fraction option). Strandedness (-s option) was specified as per the library (sample) being processed. The featureCounts annotation saf annotation file was constructed by intersecting the gene annotations (Ensembl 101) and repeatmasker file (hg38.fa.out Dec 2013 – RepeatMasker open-4.0.5 – Repeat Library 20140131), keeping intronic and intergenic TEs in R. The final gene and intergenic/intronic TE saf file was used for quantification. For each TE sub-family, its enrichment was calculated as described in Shao and Wang, 2020 Genome Research:

$$obs/exp = \frac{\frac{Number\ of\ TE\ subfamilies > 1cpm}{Number\ of\ TEs > 1cpm}}{\frac{Number\ of\ TE\ subfamilies}{Number\ of\ TEs}}$$

The observed frequency of TEs belonging to a family in all candidates divided by the expected frequency of TEs belonging to this family in genomic regions that do not overlap with protein-coding genes. To calculate the observed frequency of TEs, only the ones above 1 count per million (CPM) were considered. The pipeline and supporting files used can be accessed through the following link: https://github.com/huishenlab/TE_quantification_pipeline.git. BAMs were coordinate sorted and indexed using samtools before being used as input to the TE quantification pipeline described above. Correlations to bulk total RNA-seq were performed at the pseudobulk level and TEs found in at least 10% of cells were kept. Linear models were fit to the log(CPM+1) using the lm() function to derive adjusted $r^2$ values and plotted using ggplot2. Fallopian tube epithelium TE expression was calculated as described above, keeping TEs that were expressed (CPM > 1) in at least 10% of cells, converted to a SingleCellExperiment object, and markers were found using scran (v1.32.0) findMarkers() with the Wilcoxon Rank Sum test. Significant TE markers were filtered to those with an adjusted p-value < 0.05 and a summary AUC >= 0.9. These were further filtered to intergenic LINE, SINE, and LTRs using the annotation file described above.

Heatmaps were plotted using ComplexHeatmap and dotplots were plotted using ggplot2. Transcription factor binding enrichment was performed using FIMO (v5.5.7) for ciliated TE markers only, using a slightly relaxed set of criteria (adj. p-value < 0.05 and summary AUC >= 0.8).

**TE-derived transcript reconstruction**

TE-derived transcripts were assembled and quantified using TE-derived Promotor finder 3 (TEProf3 - https://github.com/Yonghao-Holden/TEProf3), with modifications. Briefly, resources/annotation file generation were modified to allow for use with Ensembl 101 annotations. Further, TE-derived transcript quantification using stringtie2 was modified to use the input library strandedness (e.g., reverse stranded for STORM-seq). Possible TEs (N = 4,776,620) were filtered to those that overlapped distal or proximal K-562 GRO-cap/PRO-cap TREs from PINTS (https://pints.yulab.org/; n = 37655) to restrict to expressed TEs within K-562. Initial assembly of TE-derived transcripts was performed at the pseudobulk level for K-562 STORM-seq data using -am 1, -at 40, -fm 1, -ft 8, -tt 40, -ql 150, and -ki. The assembled, filtered, TE-derived transcript GTF was then quantified at the single cell level using the –guided option in TEProf3. Bulk, total RNA-seq was also quantified using this GTF. Single-cell TE-derived transcripts were kept if expressed in at least 10% of cells and bulk-level TE-derived transcripts were filtered using recommended filters from TEProf3. Comparison to previously reported TE-derived transcript candidates in K-562 (Shah et al., 2023 Nature Genetics) was filtered to those that had the reported TE genomic location overlapping the TREs described above (35/49 candidates). Heatmaps were plotted using ComplexHeatmap (v2.20.0). Novel candidate TE-derived transcripts were filtered to those that were expressed in 1-50%, 50-80%, and 80-100% of K-562 STORM-seq cells (TPM > 0) to characterize the heterogeneity of these TE-derived transcripts (**Supplemental Table 3**). LTR1A2-PURPL and annotated PURPL isoforms were stratified using featureCounts -s 2, -p, -g transcript_id, -R BAM -a combined TE transcript and gene annotation GTF, -J, -B, -O, --largestOverlap. The reads mapping to LTR1A2-PURPL or annotated PURPL isoforms were split and plotted using Gviz in R. Minor modifications to Gviz were made to allow for the sashimi tracks to be a relative percent of reads from the combined read depth of LTR1A2-PURPL and annotated PURPL isoforms (https://github.com/biobenkj/Gviz).

**Mapping and gene expression quantification of FTE STORM-seq libraries**

Cryopreserved fallopian tube epithelium (FTE) data were first trimmed using TrimGalore (v0.6.3) with default parameters and with the following modifications: 1) --trim-n, 2) --length 36, 3) --paired, 4) --clip_R2 3, and 5) --fastqc. Trimmed FASTQ files were aligned to GRCh38 (Ensembl 102) with salmon (v1.3.0) using default parameters with the following modifications: 1) library orientation to ISR, 2) turning on the seqBias, gcBias, and posBias options due to the random priming approach in STORM, and 3) turning on the softclip option in selective-alignment mode. Gene expression was quantified using salmon and Ensembl gene annotations corresponding to the respective assembly release. Transcript counts were imported using the import_plate_txis() function in velocessor (v0.15.27 - https://github.com/trichelab/velocessor) in R (v4.0), constructing a SingleCellExperiment object. For RNA velocity inference, the FTE samples were mapped to a spliced/unspliced index generated by eisaR (v1.2.0) and salmon (v1.3.0) using Ensembl 102 assembly and annotations. Briefly, due to the full-length protocol of STORM-seq, we set featureType to extract “spliced” and “unspliced” transcript ranges at the getFeatureRanges step. All other steps were followed as described in the eisaR vignette for generating spliced/unspliced transcript files for indexing by salmon. After importing, transcript-level counts were collapsed to gene level. Transcripts per million (TPM) computed by salmon were also transformed using the izar_transform() function in velocessor as previously described. The SingleCellExperiment object containing the spliced and unspliced gene-level counts were exported as an h5ad file using zellkonvertor (v1.0.3) for import using scvelo. Quality control and normalization were performed using the scran (v1.16.0) and scater (v1.16.0) packages in R. Briefly, low quality cells were removed using the perCellQCMetrics() and quickPerCellQC() functions in scater with default parameters. Library size normalization was done using the pooling and deconvolution approach implemented in scater.

**Single-cell clustering in primary fallopian tube epithelium**

Following library size normalization, the top 10% highly variable genes across filtered cells were retained and embedded using the runPCA() function in scater, setting *ncomponents* to 20. The number of principal components to retain were estimated using the scater function getClusteredPCs(). In total 7 PCs were retained for patient 1 (high-depth sequencing) and 15 PCs were retained for patient 2 and downstream clustering. Cluster analysis was performed using the retained PCs and building a shared nearest neighbor graph with the buildSNNGraph() function in scater. Clusters were identified using the cluster_walktrap() function in igraph (v1.2.6). Both t-distributed Stochastic Neighbor Embedding (t-SNE) and Uniform Manifold and Projection (UMAP) 2-D embedding and projections were used for visualization of the data. To compute t-SNE embedding and projections, the runTSNE() function in scater was used with 20 PCs for both patient samples. For computing density preserving UMAP (densMAP) embedding and projections, the densvis package (v1.00.6) and the densmap() function, setting *n_neighbors* to 15, *n_components* to 3, and metric to “euclidean”, was used with 20 PCs for both patients. Clusters inferred as described above were then colored on t-SNE and densMAP projections.

**Integration of multi-patient primary fallopian tube epithelium STORM-seq**

Integration of both patient 1 and patient 2 STORM-seq data, as well as integration with SMART-seq2 FTE data, was performed using mutual nearest neighbors as implemented in batchelor (v1.6.3) using the fastMNN() function as described in Orchestrating Single-Cell Analysis with Bioconductor. Briefly, for STORM-seq integration, multi-batch normalization was performed using the multiBatchNorm() function in batchelor, scaling towards the lowest coverage batch - in this case is patient 2. Next, feature selection was performed by averaging the multi-patient variance components using the combineVar() function in scran (v1.16.0) and keeping genes above the trend as previously described[72]. In total, 17,573 genes were retained. Next, the rescaled patient counts and selected features were passed to fastMNN() using default parameters with the following modifications: 1) k = 15 and 2) subset.row = selected features described above. The integrated data were then used to compute clusters and a 2-D t-SNE projection similar to the process described above, with the only difference being that 50 PCs were used as input to the runTSNE() function. To integrate STORM-seq data with previously published SMART-seq2 FTE data, the process described above was repeated for integration, treating batches as 1) patient 1 STORM-seq, 2) patient 2 STORM-seq, 3) SMART-seq2 FTE data.

**Cell type annotation in primary fallopian tube epithelium**

Cell types were defined using the clusters inferred as described above, and both known cell type markers and inferred marker genes.

**Trajectory inference of primary fallopian tube epithelium**

Single-cell experiment objects were exported as an h5ad file using zellkonverter (v1.0.3). Exported h5ad files were imported into python (v3.8.5) using scanpy (v1.7.1) and scvelo (v0.2.3), converting to an AnnData object. The clusters containing immune cells were excluded for velocity inference. Each donor was processed separately, but using the same parameters for initial preprocessing and model fitting. Briefly, spliced and unspliced counts were filtered and normalized using the filter_and_normalize() in scvelo with the following modifications: 1) min_shared_counts=20 and 2) n_top_genes=1000. Next, first- and second-order moments were computed using the moments() function with the following modifications: 1) n_pcs=5 and 2) n_neighbors=15. The number of PCs and neighbors chosen for computing moments was based upon the variance explained for initial embedding of individual patients as well as consistency of nearest neighbors chosen for t-SNE, densMAP, and MNN integration. Full splicing kinetics were recovered using recover_dynamics() for inference of latent time. Next, the velocities were computed using the stochastic model within the velocity() function. The CytoTRACE pseudotime inference was performed using 10000 genes as input as implemented in CellRank (v1.3.1). Driver gene inference for lineage commitment was performed using the above RNA velocity approach and n_top_genes=10000. Principal curves were calculated using slingshot (v2.12.0) in R. Briefly, clusters were annotated based on marker gene classification to UCFP, Secretory, Ciliated, Ciliated intermediate, Secretory intermediate, Immune, and Branch across donors. The immune populations were removed from this analysis to investigate trajectories to ciliated and secretory cells, specifically. Next, getLineages() was used to infer principal curves and pseudotime within PCA space for each donor, individually. Curves were extracted and pseudotime was rescaled to 0-1 range. Curves and pseudotime were plotted using ggplot2.

**CycIF image acquisition and analysis of whole tissue sections of primary human fallopian tube**

CellDive CyCIF images were acquired using manufacturer recommended sample preparation and CellDive validated antibody concentrations from Cell Signaling Technology (NaKATPase – AF488, D4Y7E clone; PanCK – AF488, C11+E6S1S clones; CD31 – AF750, 89C2 clone; DAPI for nuclear staining). Full resolution OME TIFFs of CellDIVE images (one multipage tiff per round of staining) were imported into QuPath (v5.1) for analysis (Bankhead, P., et al., 2017). All multipage tiffs were merged to generate an image containing all rounds of staining; because CellDIVE images are automatically aligned during acquisition, the Interactive Image Combiner Warpy tool (Chiaruttini, N., et al., 2022) was used to merge images from all rounds of staining without application of a transformation matrix. A training image composed of regions of interest (ROIs) from the merged image from all samples was generated to establish cell segmentation and object classification parameters. Cells were segmented by running the CellPose cyto3 model (Stringer and Pachitariu, 2024) on the NaKATPase channel and the DAPI channel acquired in the same round of staining and imaging. Random Trees object classifiers were trained to identify CD31-, PanCK-, or CD45-positive cells independently (the mean, median, min, max, and std. dev. values for the respective channels were used for training). These three classifiers were used to generate a composite classifier to identify single (CD31- or PanCK-positive, and CD45-negative) and dual-feature (CD31; PanCK-positive and CD45-negative) cells. Percentage of dual-feature cells were calculated based on the composite classifier.
