## Supplemental Figures for "Efficient profiling of total RNA in single cells with STORM-seq"

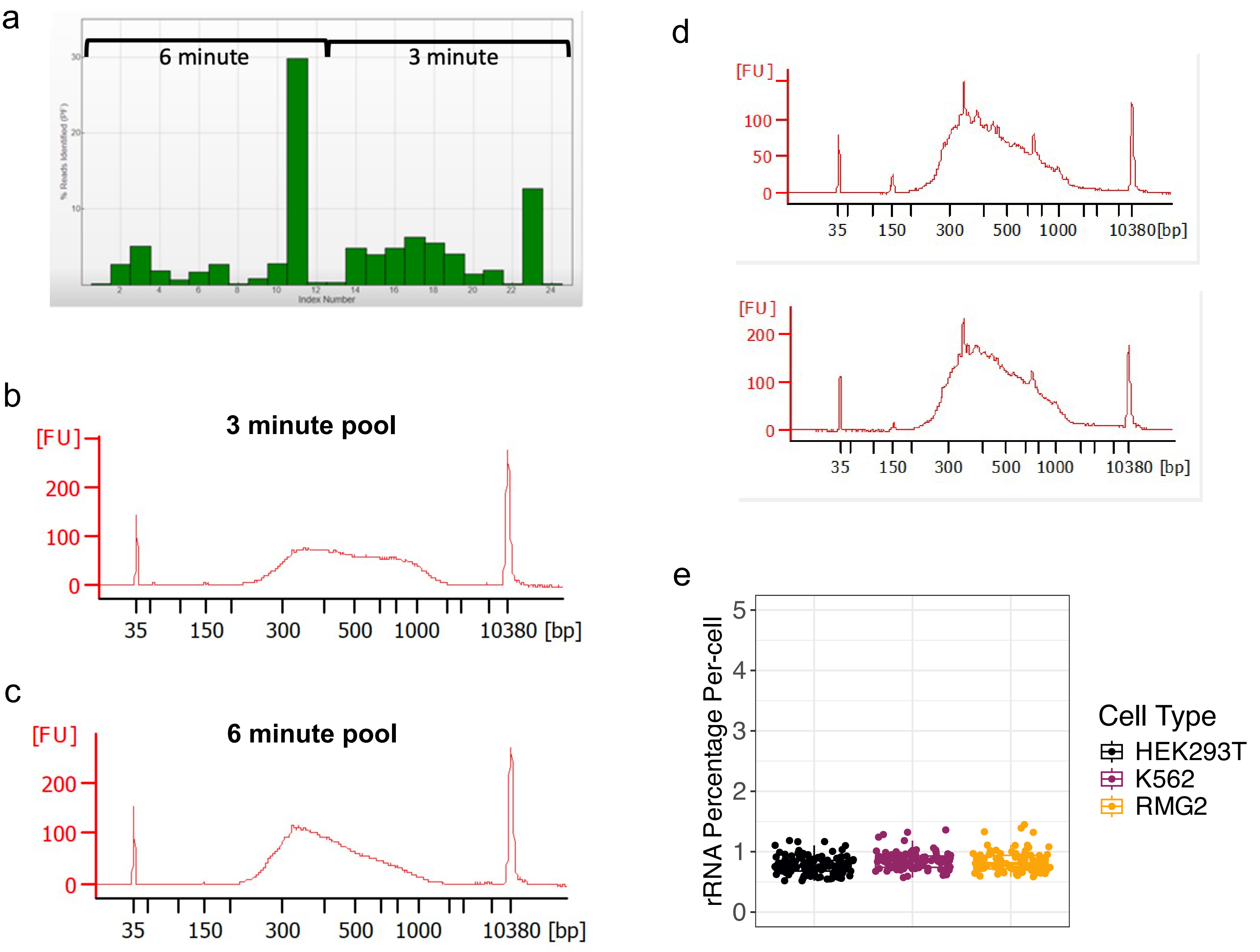


**Supplemental Figure 1. Reducing fragmentation time resulted in more even coverage (read depth) of single cells and more uniform fragmentation patterns in single-cell pools and ribosomal RNA (rRNA) is effectively depleted in STORM-seq.** a) Illumina Sequencing Analysis Viewer shows shorter fragmentation time decreases cell to cell variability. b) and c) Agilent Bioanalyzer traces for libraries prepared using 3 minute and 6 minute fragmentation times. d) Representative Bioanalyzer traces for two separate 384-cell STORM-seq pools. e) Following rRNA depletion, STORM-seq shows consistent, low total rRNA UMI count proportions per cell, across cell types, indicating effective rRNA depletion. Cells were subsampled to 100k raw reads/cell for consistency with prior analyses in Fig. 1. rRNA genes included are from Ensembl 101 gene annotations and include: rRNA, rRNA_pseudogene, and Mt_rRNA biotypes. A total of 99 out of a possible 551 rRNA-related genes were de-

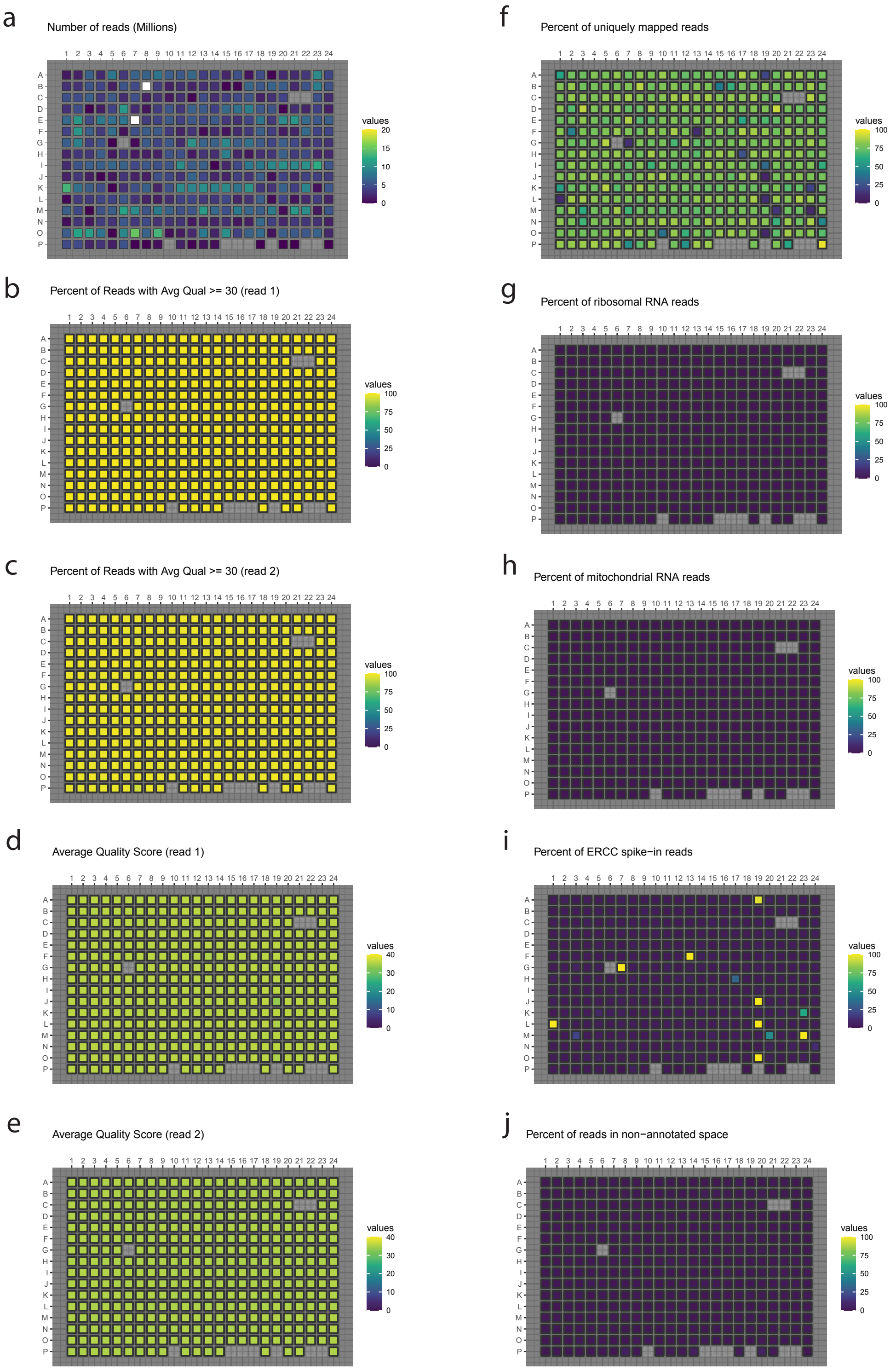


**Supplemental Figure 2. Quality metrics generated using STORMqc pipeline on a STORM-seq plate of HEK293T, K-562, and RMG-2 single cells. a)** Number of reads (millions). Empty boxes indicate no sequence data above minimum read threshold (100 reads). White squares indicate read depth beyond the upper bound of color scale (20 million reads). **b)** Percent of reads with average base quality >= 30 (read 1, Phred 33 scale) **c)** Percent of reads with average base quality >= 30 (read 2, Phred 33 scale) **d)** Average base quality score (read 1, Phred 33 scale) **e)** Average base quality score (read 2, Phred 33 scale). **f)** Percent uniquely mapped reads of total. **g)** Percent of ribosomal RNA reads **h)** Percent mitochondrial RNA reads of total. **i)** Percent ERCC spike−in reads of total. **j)** Percent of reads in non−annotated space.

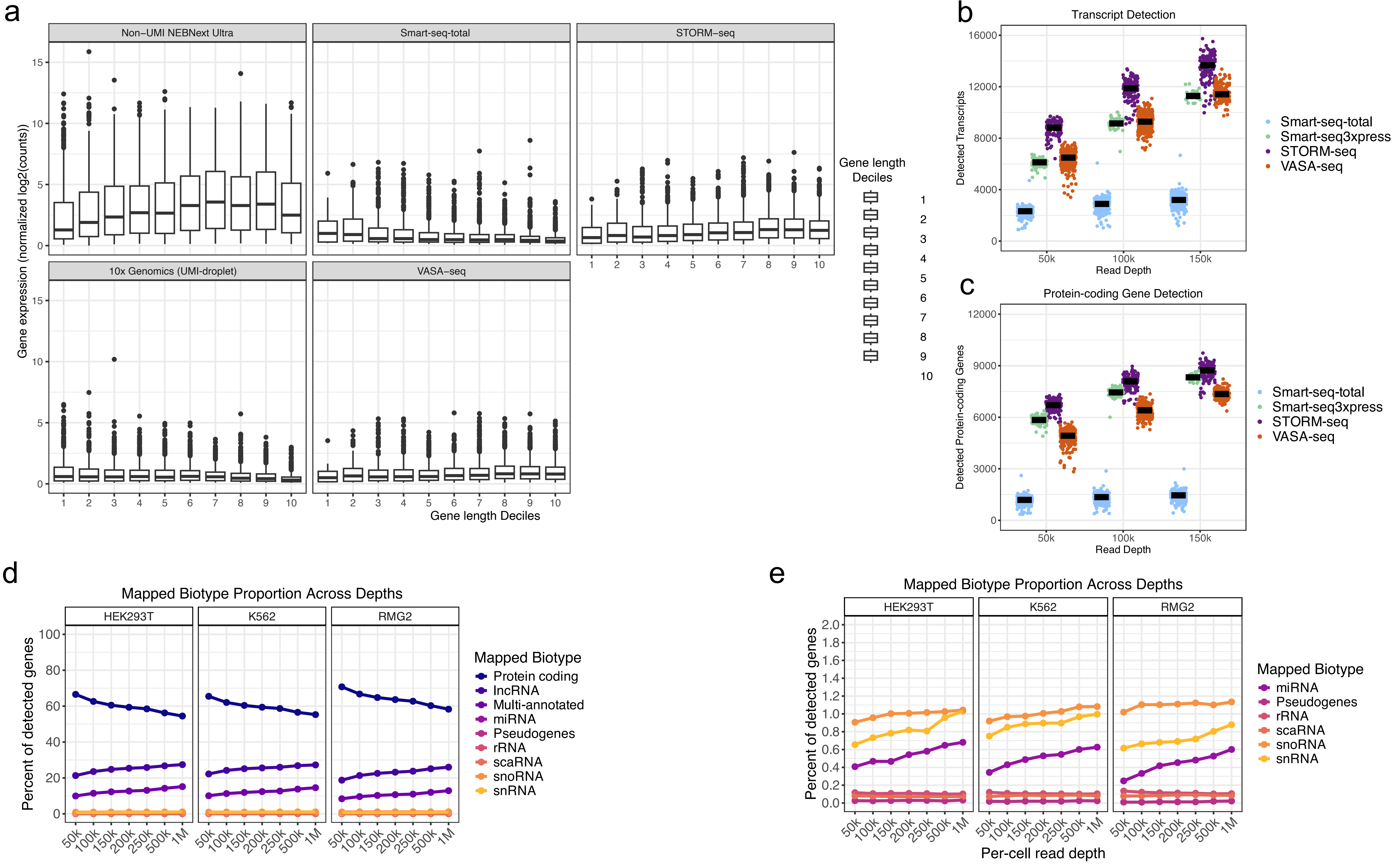


**Supplemental Figure 3. STORM-seq gene- and transcript-level detection complexity and length bias. a)** Annotated Ensembl genes were stratified into even deciles (2681 ± 1.43 genes/decile) based on their annotated lengths. Pseudobulk single-cell normalized gene expression is shown where the Non-UMI NEBNext Ultra exemplifies gene length bias with increasing gene expression as a function of gene length. The 10x Genomics pseudobulk exemplifies how end-counting protocols do not exhibit strong gene length bias (e.g., Smart-seq3xpress 5' UMI reads). STORM-seq and VASA-seq show mild gene length bias, whereas Smart-seq-total shows more bias towards shorter gene lengths. **b)** Transcript isoform detection of annotated genes in Ensembl 101 using kallisto|bustools, detection threshold minimum of at least 1 UMI count. **c)** Ensembl 101 protein-coding gene detection rates across scRNA-seq technologies, detection threshold minimum of at least 1 UMI count. All subsampling reflects the number of raw reads prior to pseudoalignment and quantification with kallisto|bustools. **d)** Gene biotype diversity as a fraction of the total detected biotypes found in Ensembl 101 across cell types and read depths (pseudobulk, UMI count > 0). **e)** Zoomed in view of low fractional abundance gene biotypes in STORM-seq.

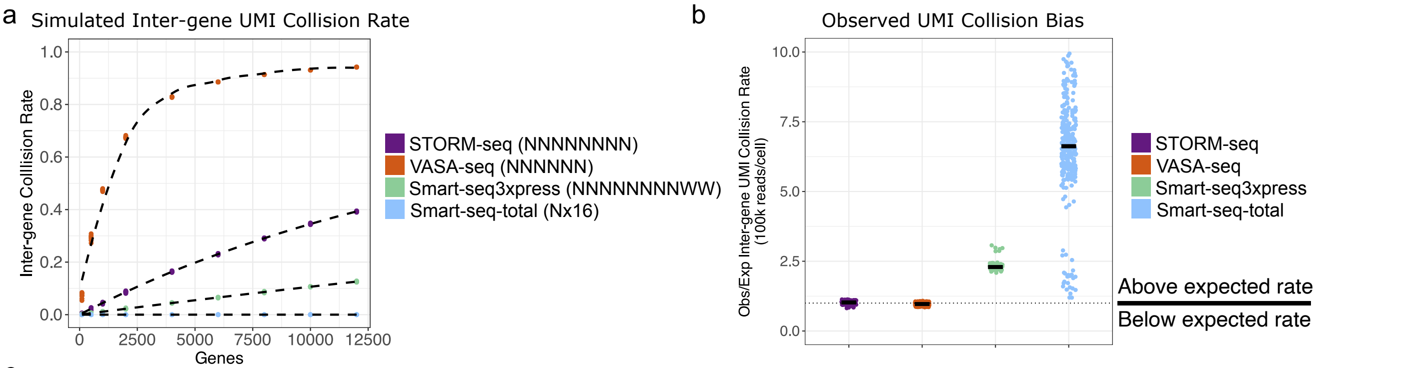


c

| Technology | UMI Motif | UMI Length | Theoretical UMI Diversity | Observed UMI Diversity* | Observed UMI Saturation* |
|---|---|---|---|---|---|
| STORM-seq | NNNNNNNN | 8 | 65536 | 65531 | 99.99% |
| VASA-seq | NNNNNN | 6 | 4096 | 4093 | 99.92% |
| Smart-seq-total | Nx16 | 16 | 4294967296 | 23131777 | 0.54% |
| Smart-seq3xpress | NNNNNNNNWW | 10 | 262144 | 430027 | 164.04% |

*Note:* *100k reads/cell

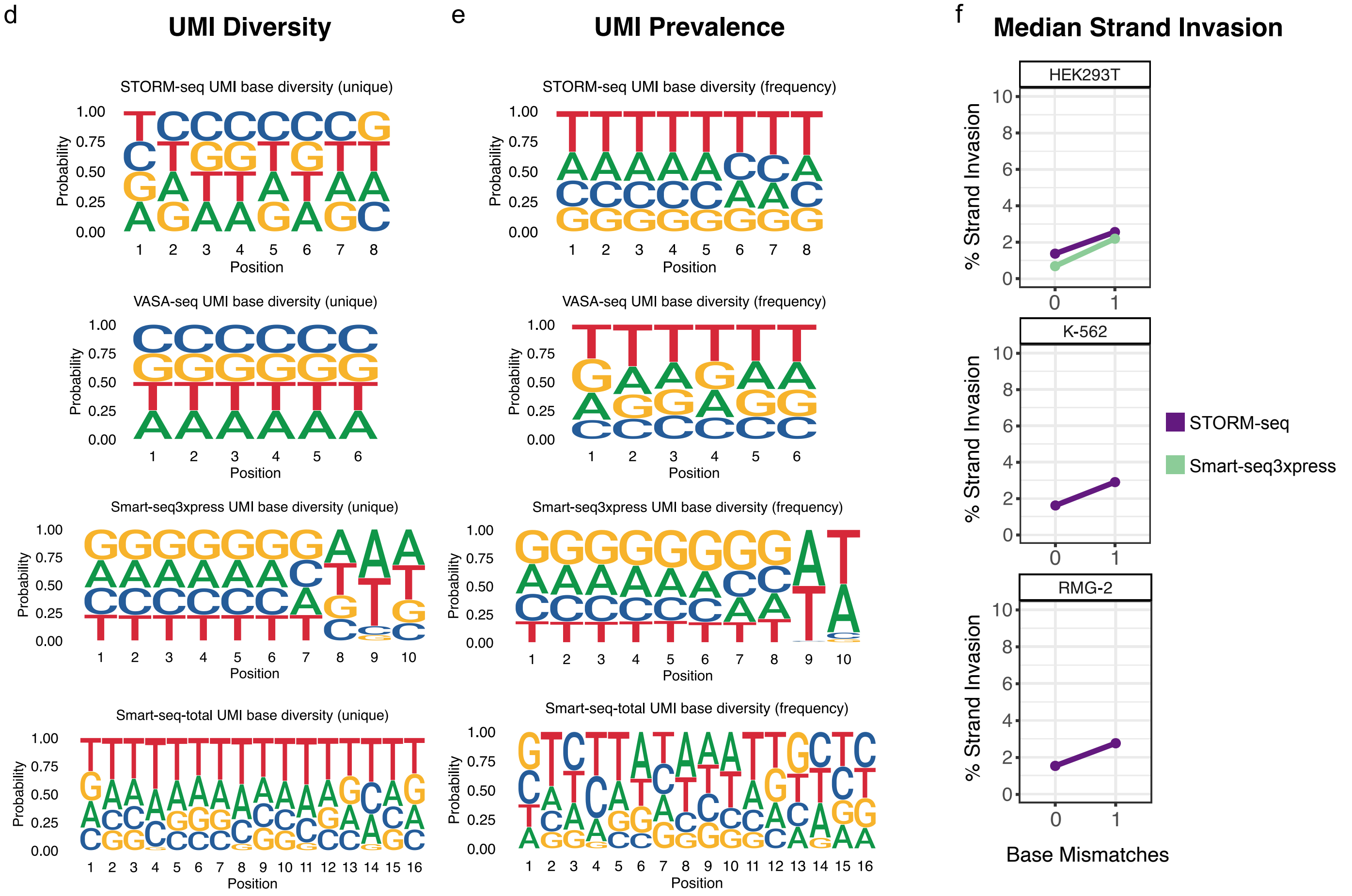


**Supplemental Figure 4. UMI motif analyses across technologies in HEK293T and strand invasion across cell types. a)** Simulated inter-gene collision rates using technology specific UMI sequences as a function of gene detection rate (proxy for read depth), to establish the expected collision rates. Simulation details and parameterizations described in the Methods. **b)** Single-cell observed/simulation expected inter-gene collision rates at a detection rate of 8000 genes/cell (expected) and 100k reads/cell (observed). **c)** Table describing UMI motif, length, theoretical, and observed diversity and saturation. **d)** Sequence logo plots of the per-base nucleotide diversity of unique, observed UMIs across single cells (diversity) by technology. **e)** Sequence logo plots of the per-base nucleotide diversity of all observed UMIs across single cells (prevalence) and technologies. **f)** Percent strand invasion allowing for zero or one mis-match in the upstream UMI match sequence. Median percentages are shown comparing STORM-seq and Smart-seq3xpress.

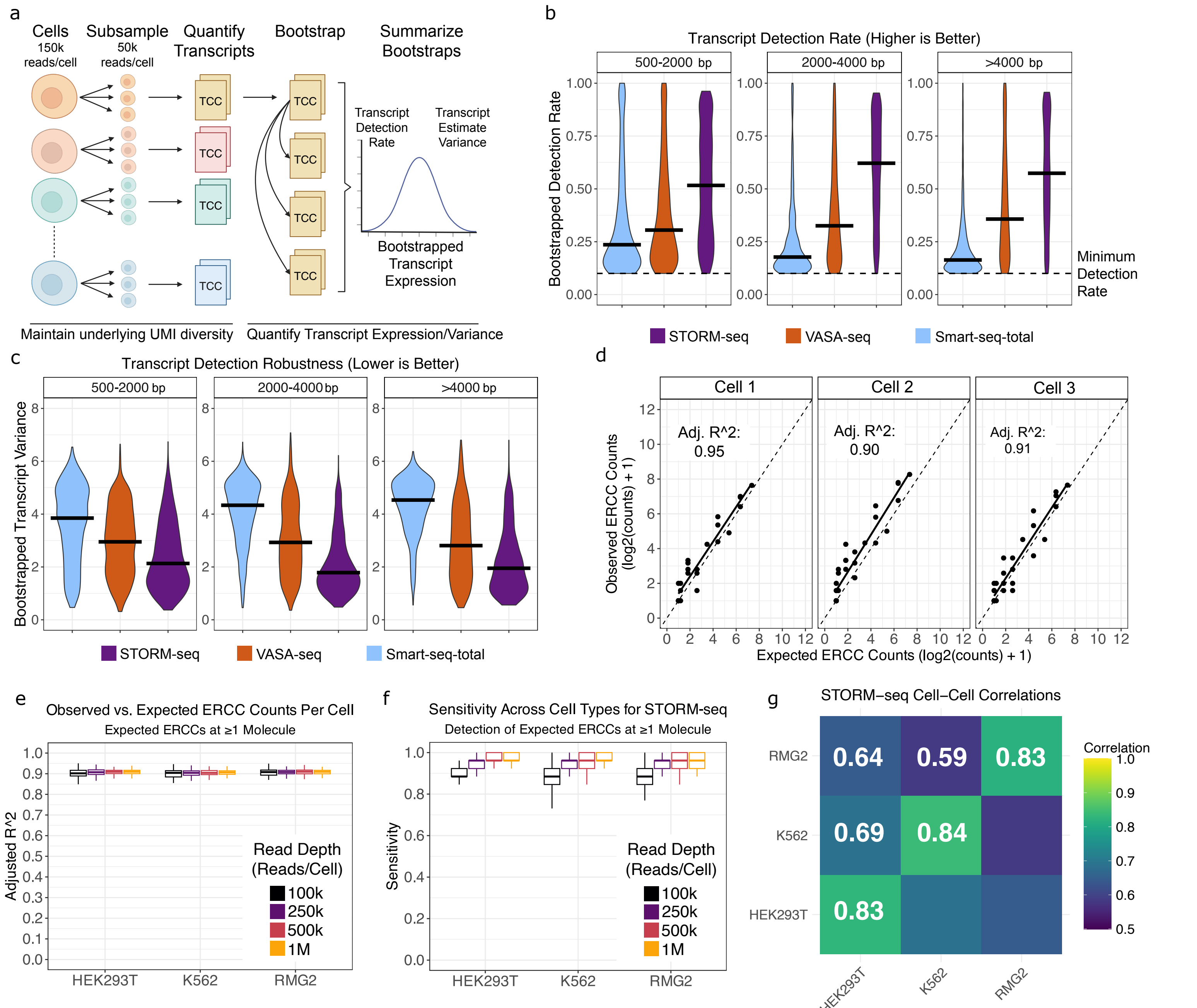


**Supplemental Figure 5. STORM-seq quantifies transcripts robustly and sensitively. a)** Schematic representation of the sampling/re-sampling scheme for panels b) and c) to simulate multiple experiments. **b)** Bootstrapped transcript detection rate (>0 UMI counts) across single cells, stratified by annotated transcript lengths. Higher is better. **c)** Bootstrapped transcript estimate variance (detection robustness), stratified by annotated transcript lengths. Lower variance estimates is better, indicating a more robust expression estimate. **d)** STORM-seq representative single cells observed/expected model fits for ERCC spike-ins. **e)** Observed/Expected model fits in STORM-seq across single cells, sequence depths, and cell types of ERCC spike-ins that are expected to have at least 1 molecule present at the 1:1M dilution. **f)** Detection sensitivity of a single molecule (ERCC) in STORM-seq data across single cells, sequence depths, and cell types. **g)** Mean cell-cell correlations of STORM-seq data (1M reads/cell). All technology comparisons were done in HEK293T cells, unless noted otherwise.

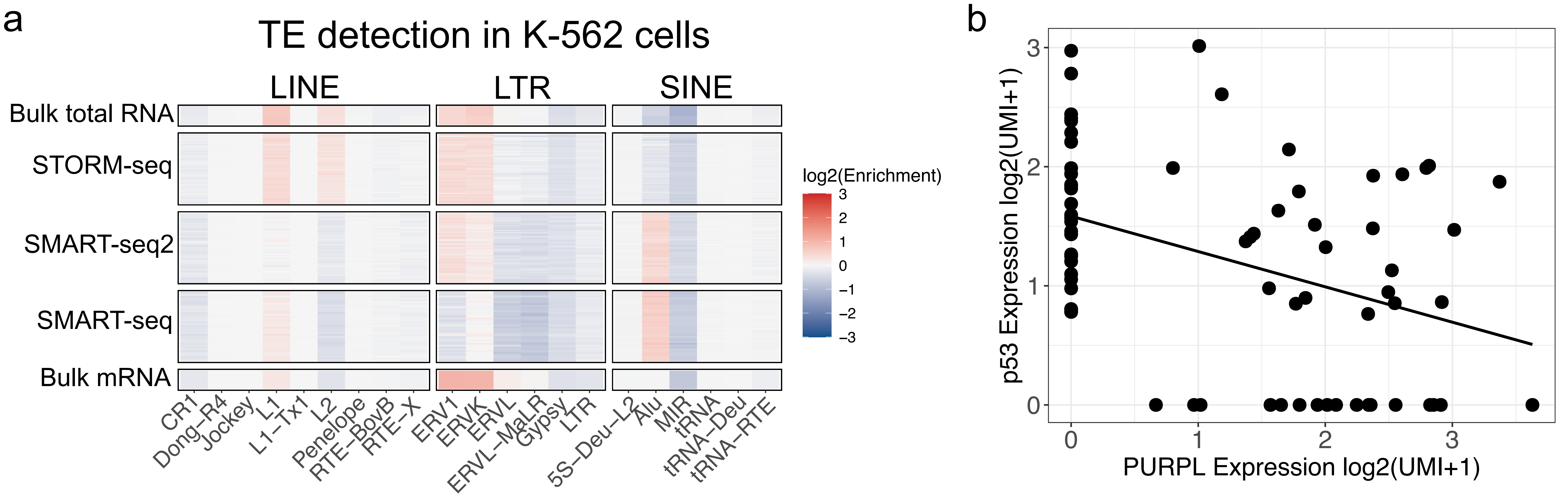


**Supplemental Figure 6. Transposable element (TE) detection in K-562 single-cells across technologies and anticorrelated *PURPL* and *p53* expression. a)** Single-cell transposable element (TE) expression representation (obs/exp) across single-cell technologies shown and bulk RNA-seq. Bulk total RNA-seq is considered the “gold standard” and bulk mRNA (oligo(dT) primed) is shown as a relative comparison. STORM-seq reconstructs bulk total RNA-seq TE profiles in single cells compared to other technologies. **b)** Expression of STORM-seq K-562 cells expressing PURPL and p53 showing expected anticorrelated expression profiles.

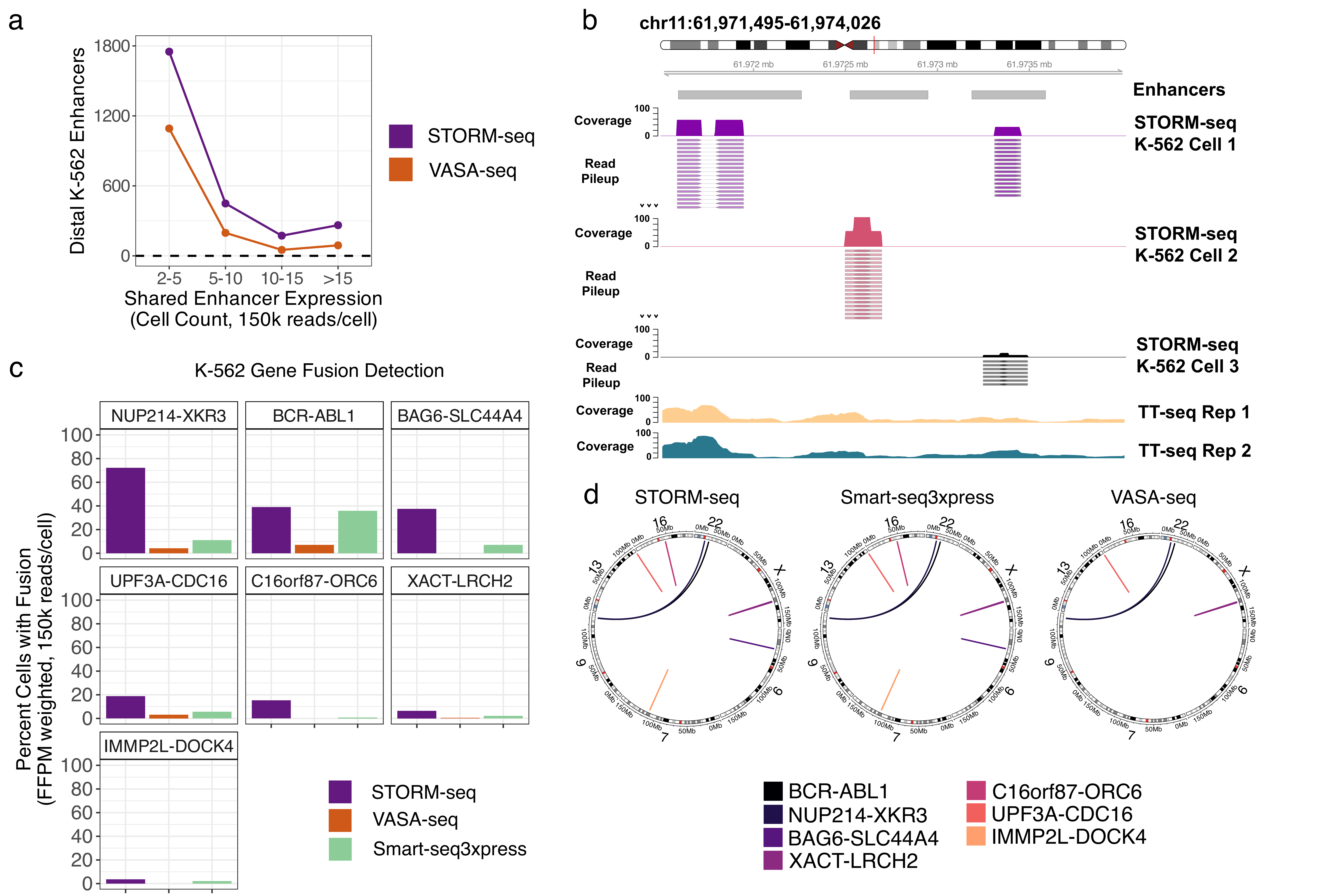


**Supplemental Figure 7. STORM-seq reconstructs cell-type specific regulatory elements and clinically relevant gene fusions in single-cells. a)** Detection rates of single-cell distal eRNAs when subsampled to 150k reads/cell shows STORM-seq identifies approximately twice as many eRNAs per cell compared to VASA-seq (left panel; black dots are median eRNA detection rates; n=100 cells STORM-seq; n=185 cells VASA-seq). Subsampling to 100 cells per technology at 150k reads/cell, STORM-seq shows more shared eRNA expression profiles compared to VASA-seq in distal K-562 enhancers (right panel). **b)** Example of single-cell differential enhancer usage in the same enhancer cluster in K-562 STORM-seq data. TT-seq data demonstrates nascent RNA signal across the annotated enhancers. Arrows indicate additional read depth not shown. **c)** Proportion of single cells with a detected, known gene fusions in K-562. Cell type proportions are weighted by respective bulk RNA-seq technology detection sensitivity to make total RNA and mRNA single-cell protocols comparable. Single cells were downsampled to 150k reads/-cell. **d)** Circos plot showing genomic alterations of known gene fusions (CCLE) in K-562. STORM-seq and Smart-seq3xpress detect known gene fusions, while VASA-seq does not.

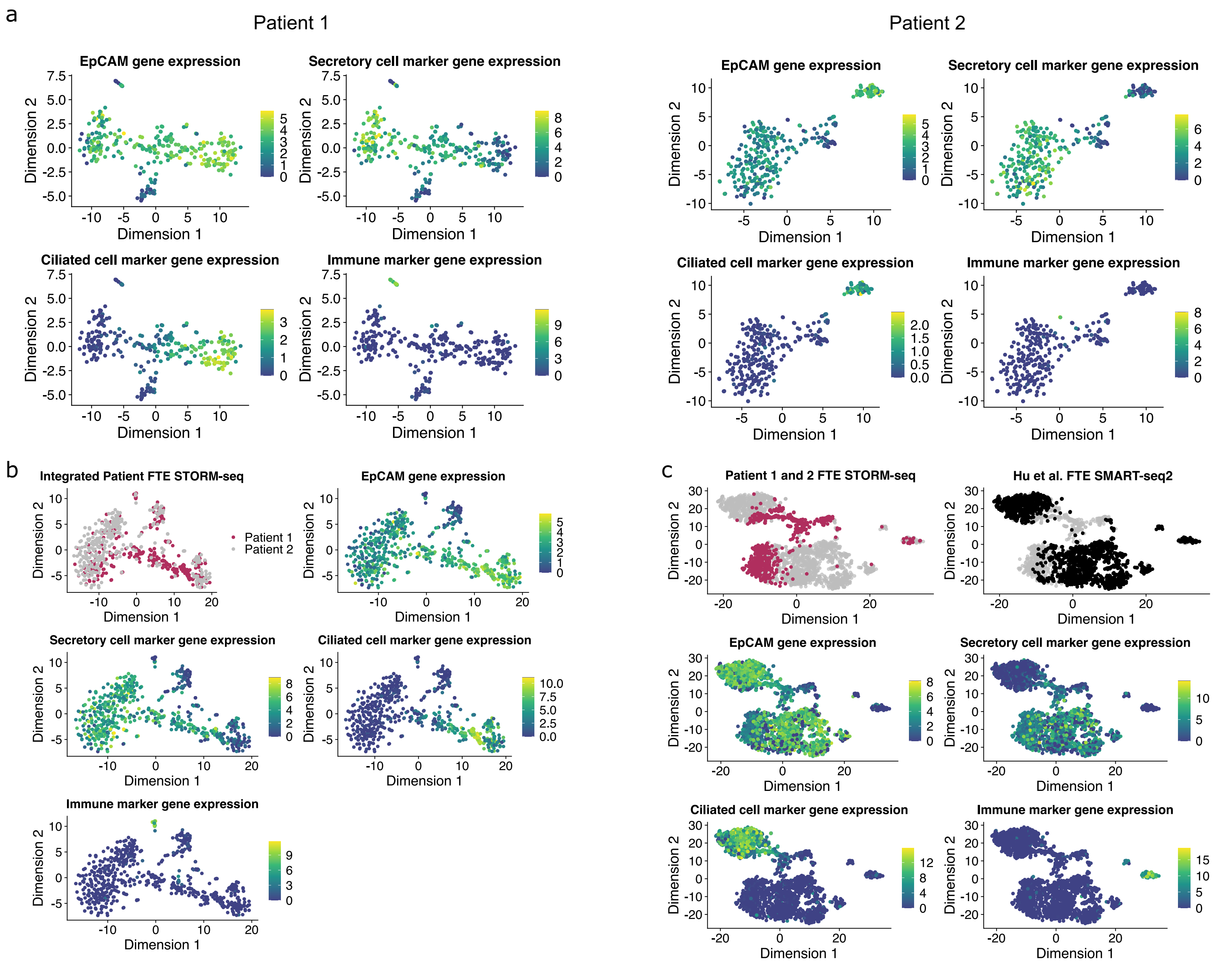


**Supplemental Figure 8. STORM-seq recovers known and intermediate cell types in primary human fallopian tube epithelium across patients and read depths.** a) Patient 1 (deep sequencing, >2M reads/cell) and patient 2 (shallow sequencing, <1M reads/cell) primary fallopian tube epithelium (FTE) were compared for recovery of known cell types within the FTE. Log2(TPM+1) is shown. Marker gene expression was summed. Secretory cell marker genes: PAX8 and KRT7. Ciliated cell marker genes: FOXJ1, CCDC17, and CCDC78. Immune cell marker genes: CD45, ITGAX (CD11c), and CD14. b) Integration of patients 1 and 2 via mutual nearest neighbors (MNN) shows that STORM-seq recovers known and intermediate cell types in FTE across patients and read depths. Log2(TPM+1) is shown. c) Integration of STORM-seq with normal FTE from SMART-seq2 (Hu et al. 2019, Cancer Cell) via MNN demonstrates recovery of known cell types in the FTE across protocols. Notably, STORM-seq recovers more intermediate cells than SMART-seq2. All embeddings shown are t-SNE. Gating strategy for cell sorting was singlet, viable (DAPI dim), EpCAM+, CD238a- cells.

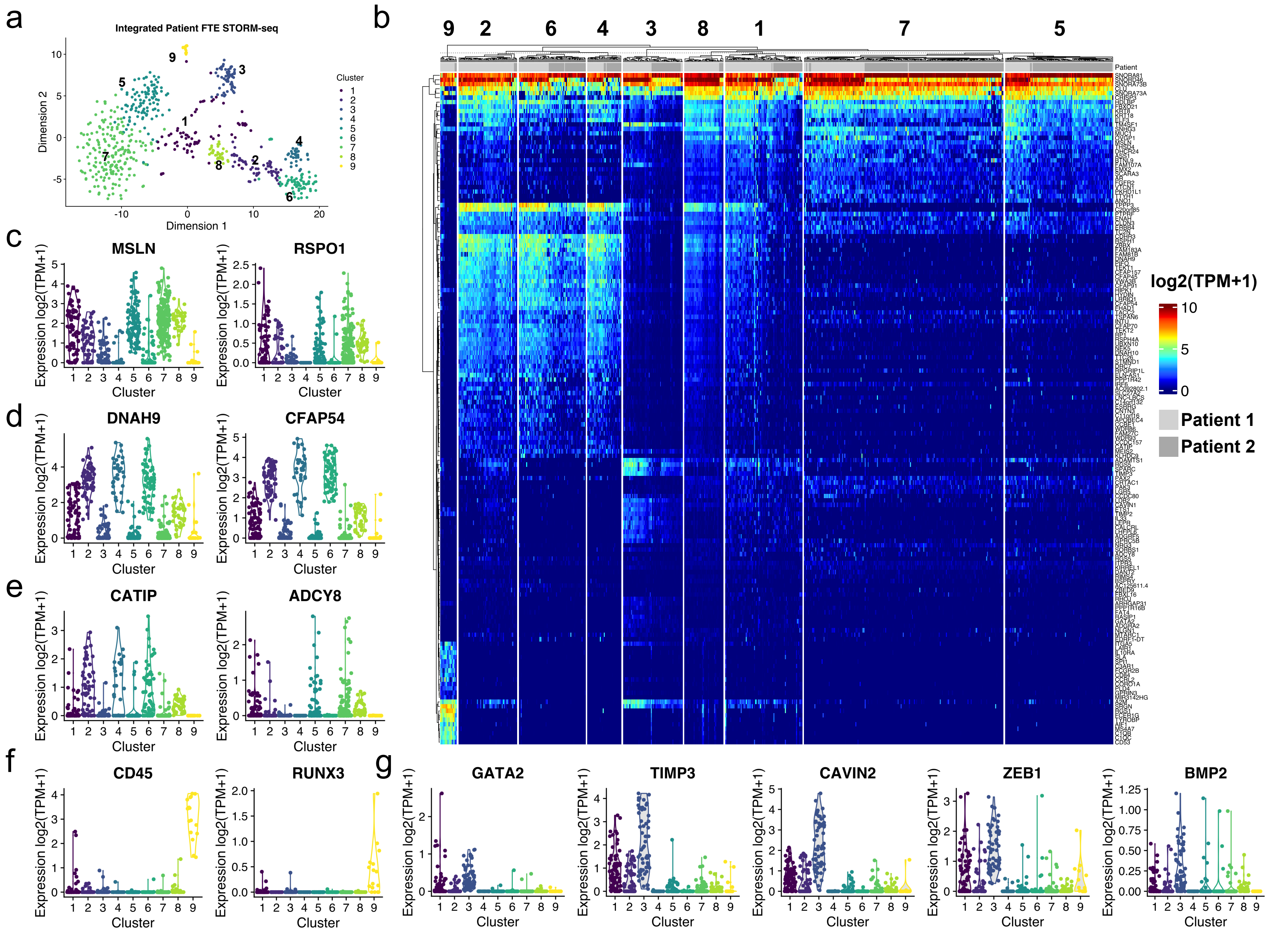

**Supplemental Figure 9. Marker gene detection in the primary FTE identifies novel gene programs for early, intermediate, and terminally differentiated cell types.** a) MNN-integration of patients 1 and 2 identifies distinct clusters of UCFP (cluster 3), “intermediate” (cluster 8), ciliated intermediate cells (cluster 2), secretory intermediate cells (cluster 1), and multiple subpopulations within secretory (clusters 5 and 7) and ciliated (clusters 4 and 6) terminally differentiated cells. Further, a unique population of EpCAM+, CD45 expressing cells was also identified (cluster 9). b) Heatmap of the top 20 marker genes per cluster. Top marker genes were identified using pairwise Wilcoxon rank sum tests, and keeping significant ($q < 0.05$) genes that were discriminatory for their cluster based on area under the curve (AUC). c) Example marker gene expression for secretory cells (clusters 5 and 7). d) Example marker gene expression for ciliated cells (clusters 4 and 6). e) Example marker gene expression for “intermediate” cells (cluster 8). f) Example marker gene expression for “immune” population (cluster 9). g) Example marker gene expression for cluster 3 (UCFP, progenitor population). All expression values shown are log2(TPM+1).

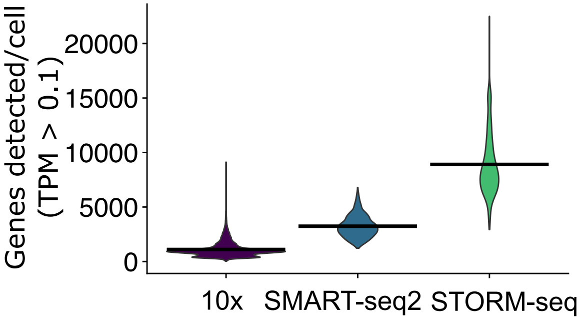


**Supplemental Figure 10. Gene detection across technologies in human FTE.** Gene detection rates across single cells in primary human fallopian tube epithelium across technologies. Mean gene detection rates: 10x genomics – 1102 genes/cell; SMART-seq2 – 3238 genes/cell; STORM-seq – 8906 genes/cell. 10x Genomics data: Dinh et al. 2021 Cell Reports; SMART-seq2: Hu et al. 2020 Cancer Cell.

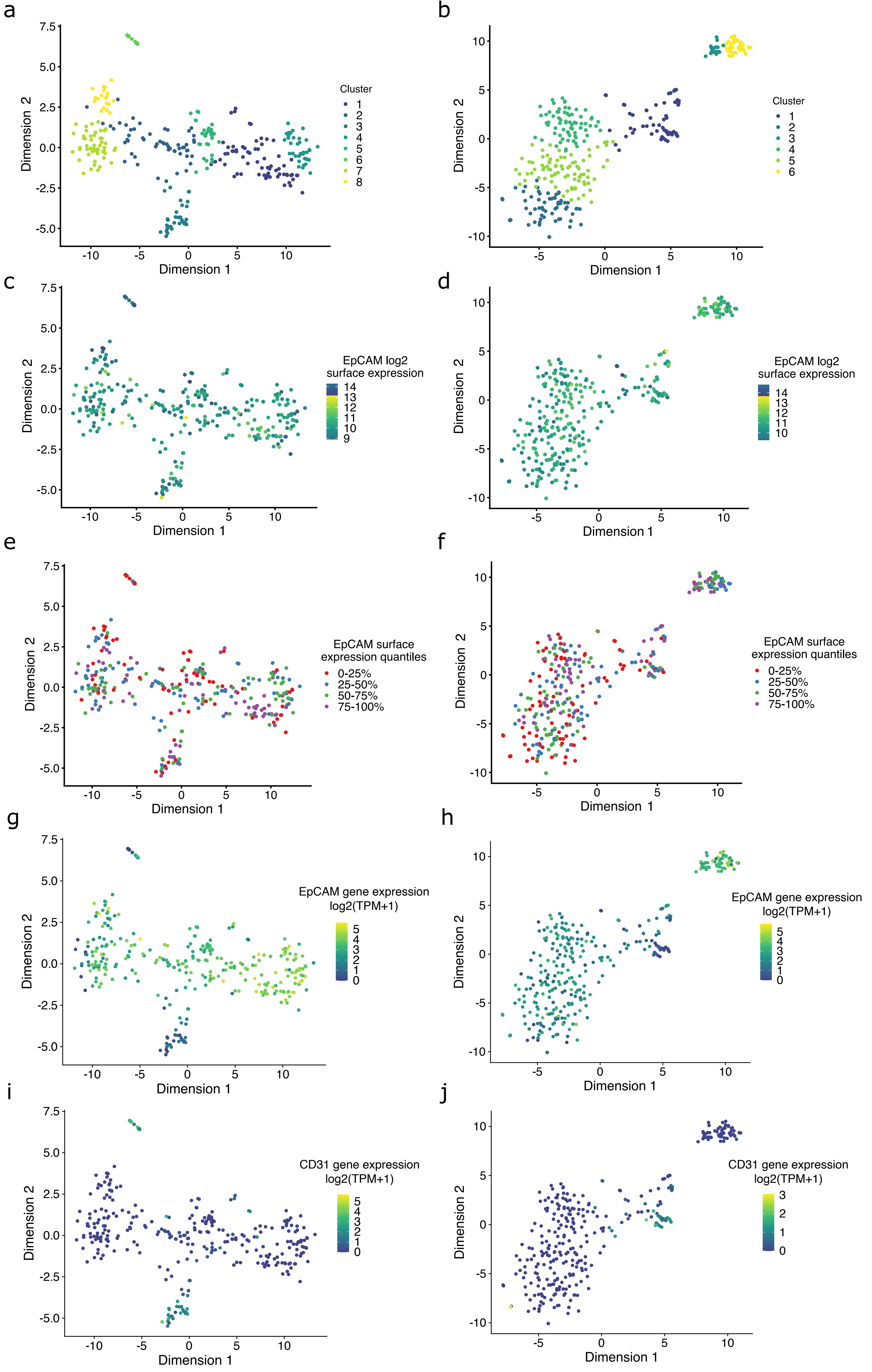


**Supplemental Figure 11. UCFP "dual-feature" epithelial and endothelial cell surface and gene expression.** a) Patient 1 and b) patient 2 detection of cell populations using STORM-seq. t-SNE embeddings shown. c) Patient 1 and d) patient 2 log2 cell surface expression of EpCAM across cell types and clusters. e) Patient 1 and f) patient 2 quantiles of EpCAM surface expression further supports the even distribution of EpCAM cell surface expression across clusters and cell types. g) Patient 1 and h) patient 2 EpCAM gene expression. i) Patient 1 and j) patient 2 CD31/PECAM1 gene expression.

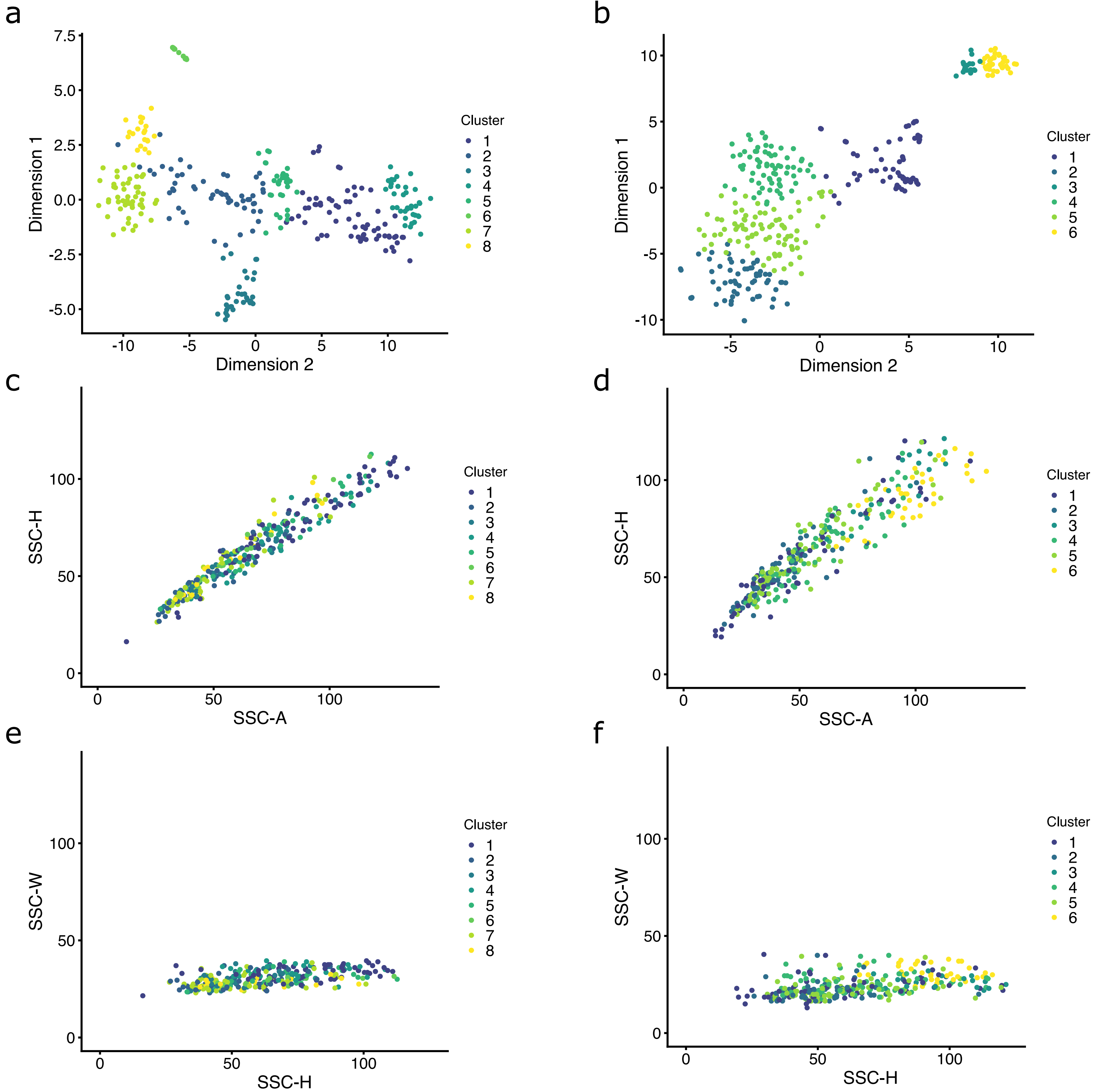


**Supplemental Figure 12. Cluster and cytometry-based detection of doublets in primary human fallopian tube epithelium.** a) Patient 1 and b) patient 2 detection of cell populations using STORM-seq. t-SNE embeddings shown. c and e) Index sorted patient 1 and d and f) patient 2 side-scatter area vs side-scatter height plots and side-scatter height vs side-scatter width plots colored by cluster demonstrate no doublets were detected. Expected doublets would occur in the lower right quadrants of c and d) and the upper quadrant of e and f). We observe tight population clustering with no individual cells/clusters forming doublets in the expected quadrants.

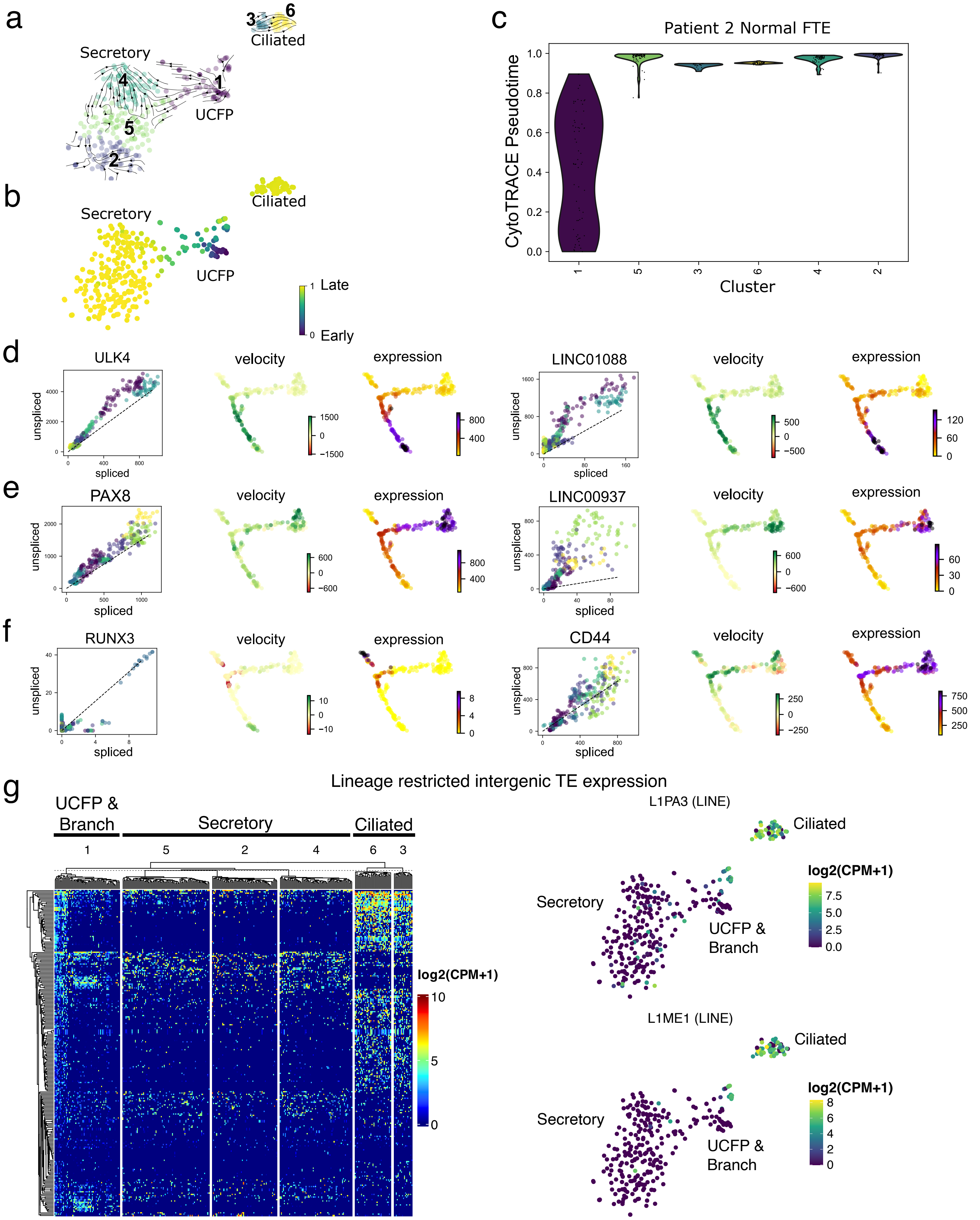


**Supplemental Figure 13. Patient 2 RNA velocity and CytoTRACE pseudotime show similar trajectory from a bipotent UCFP population of cells towards terminally differentiated cell types and phase portrait example driver genes from patient 1. a)** RNA velocity from the UCFP population of cells (cluster 1) towards ciliated (clusters 3 and 6) and secretory (clusters 2, 4, and 5). **b)** CytoTRACE pseudotime supports the inferred RNA velocity with early cells being the UCFP cluster and late cells are the terminally differentiated populations. **c)** CytoTRACE pseudotime ordering of single cells, stratified by cluster. t-SNE embeddings shown. **d)** Ciliated and **e)** secretory cell phase portraits of RNA velocity driver genes indicate *ULK4* (ciliated), *PAX8* (secretory), and long intergenic non-coding RNAs as critical for differentiation trajectories. **f)** *RUNX3* shows low velocity and does not support a dedifferentiation trajectory. *CD44* is a proposed marker for stem-like cells in human FTE. Colors in the spliced/unspliced phase portrait are colored by cluster as in A). Green shows positive velocity and red negative velocity. Expression is modeled by log transformed gene counts. RNA velocity, CytoTRACE pseudotime, and latent time were calculated using scvelo. densMAP embeddings shown for patient 1. **g)** Representative cell-type specific intergenic transposable element (TE) expression from patient 2 demonstrates lineage restricted TE expression, with the same locus-level LINE expression within the ciliated cell lineage also found in patient 1. t-SNE embeddings shown for patient 2.

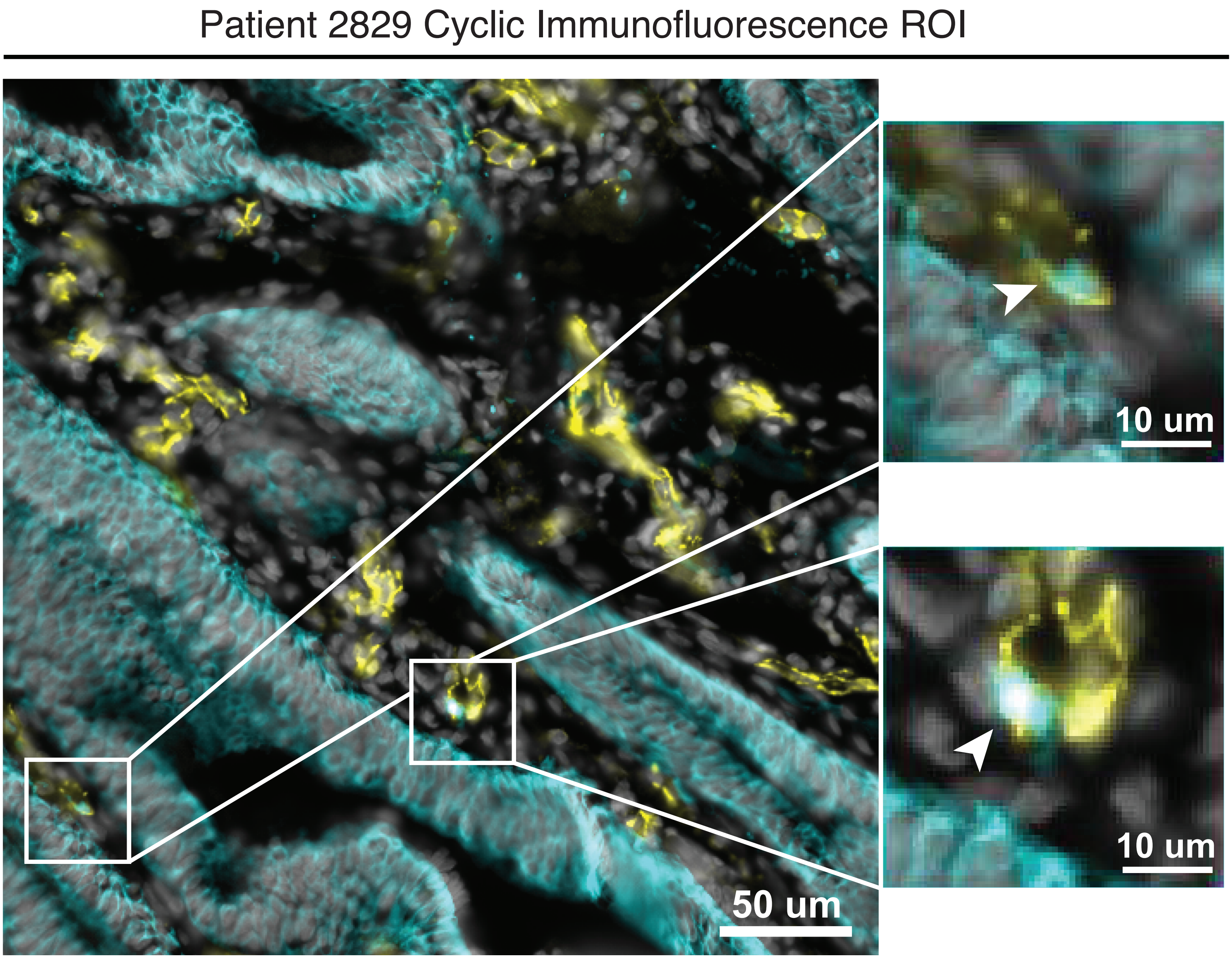


**Supplemental Figure 14. Patient 2829 example "dual-feature" cells from cyclic immunofluorescence.** Separate regions of interest (ROI) with zoomed insets to show representative simultaneous expression of epithelial (PanCK) and non-epithelial (CD31) cells.

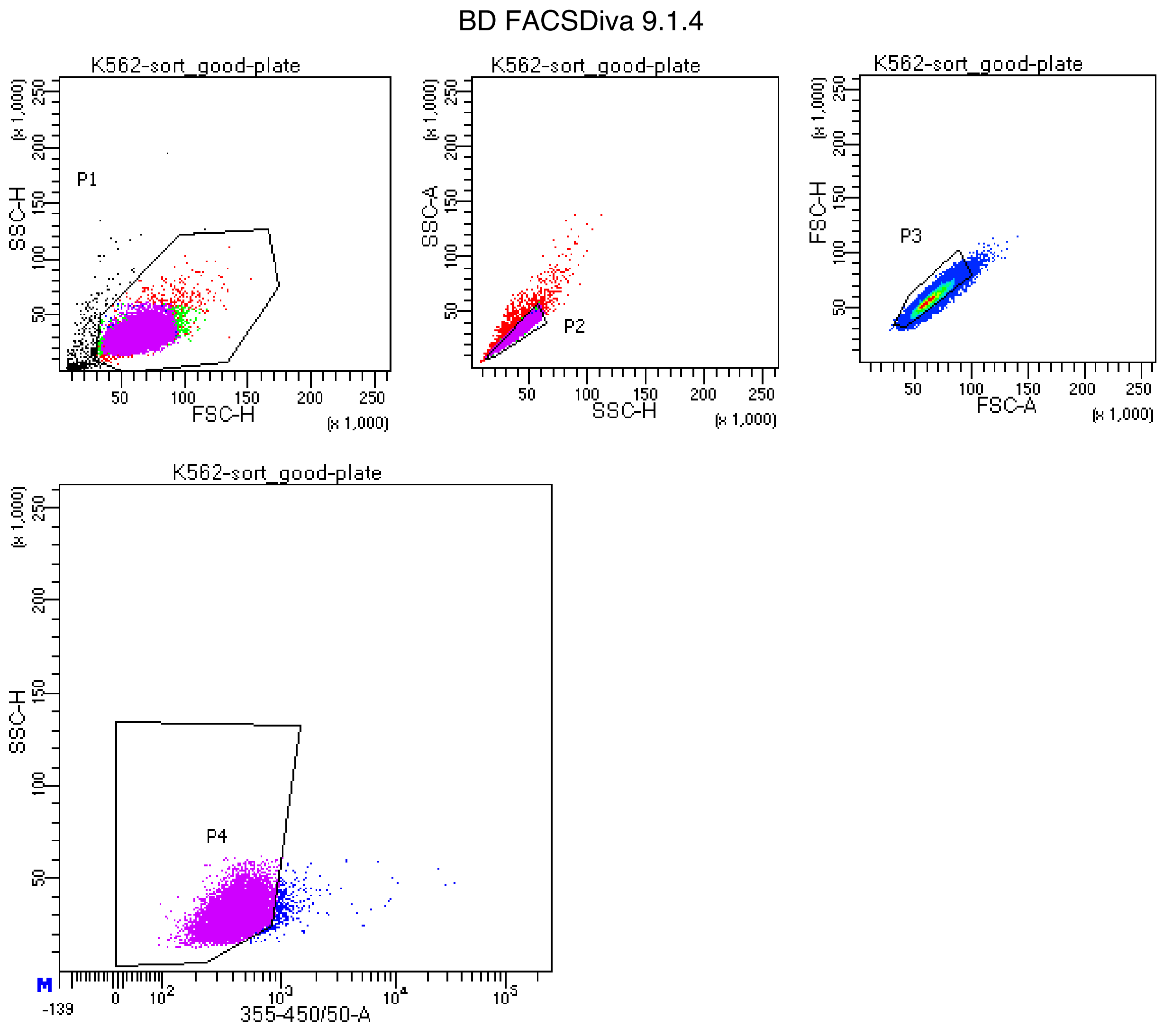


Tube: sort_good-plate

| Population | #Events | %Parent | %Total |
|---|---|---|---|
| All Events | 24,169 | #### | 100.0 |
| P1 | 22,967 | 95.0 | 95.0 |
| P2 | 20,308 | 88.4 | 84.0 |
| P3 | 19,560 | 96.3 | 80.9 |
| P4 | 19,153 | 97.9 | 79.2 |

**Supplemental Figure 15. Representative K-562 flow cytometry gating scheme for STORM-seq and VASA-seq libraries.** Gating strategy gating on cells (P1), single cells (P2), single cells (P3), and viablity (P4 - DAPI).

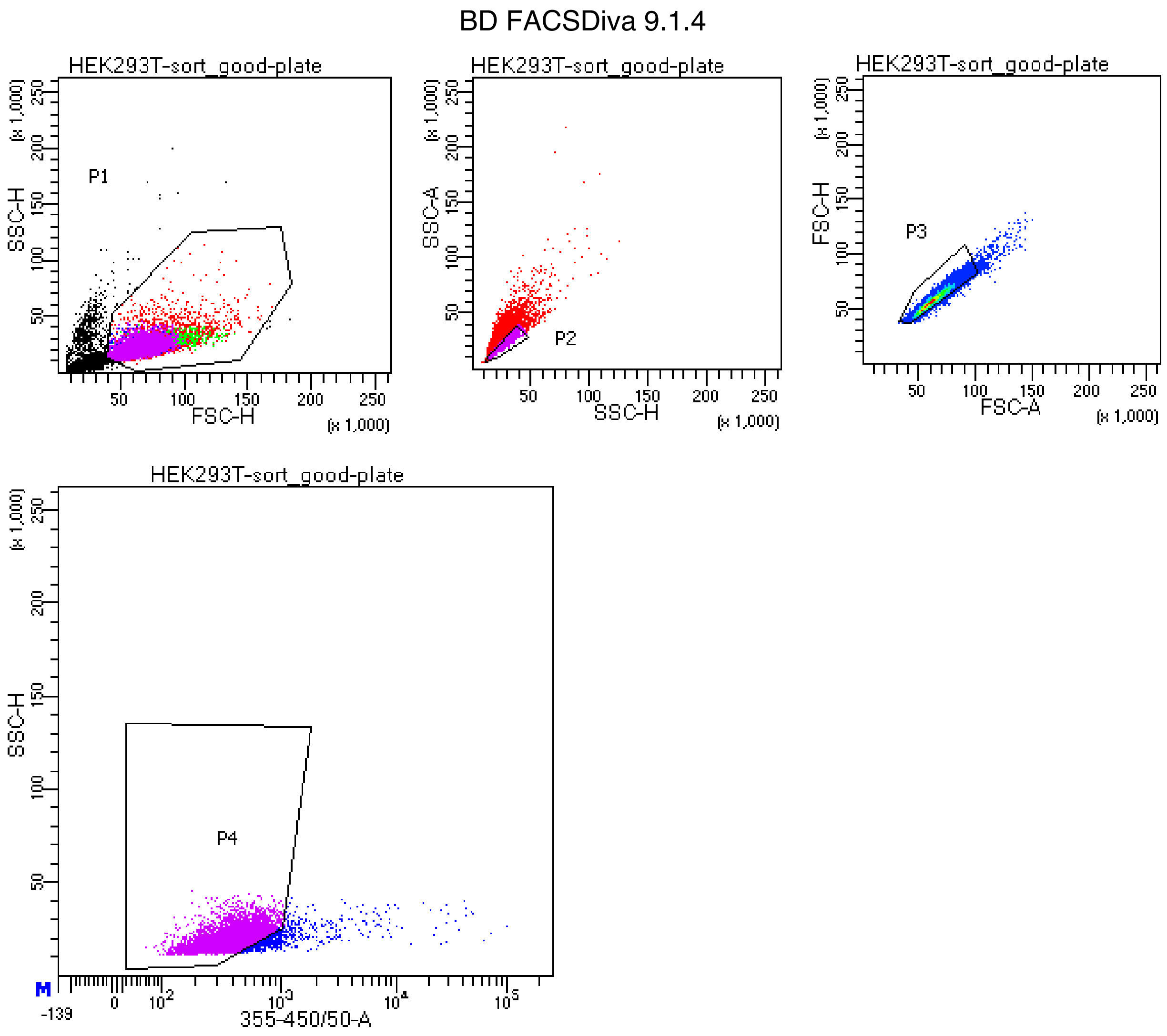


Tube: sort_good-plate

| Population | #Events | %Parent | %Total |
|---|---|---|---|
| All Events | 29,988 | #### | 100.0 |
| P1 | 24,308 | 81.1 | 81.1 |
| P2 | 17,415 | 71.6 | 58.1 |
| P3 | 17,072 | 98.0 | 56.9 |
| P4 | 16,015 | 93.8 | 53.4 |

**Supplemental Figure 16. Representative HEK293T flow cytometry gating scheme for STORM-seq and VASA-seq libraries.** Gating strategy gating on cells (P1), single cells (P2), single cells (P3), and viablity (P4 - DAPI).

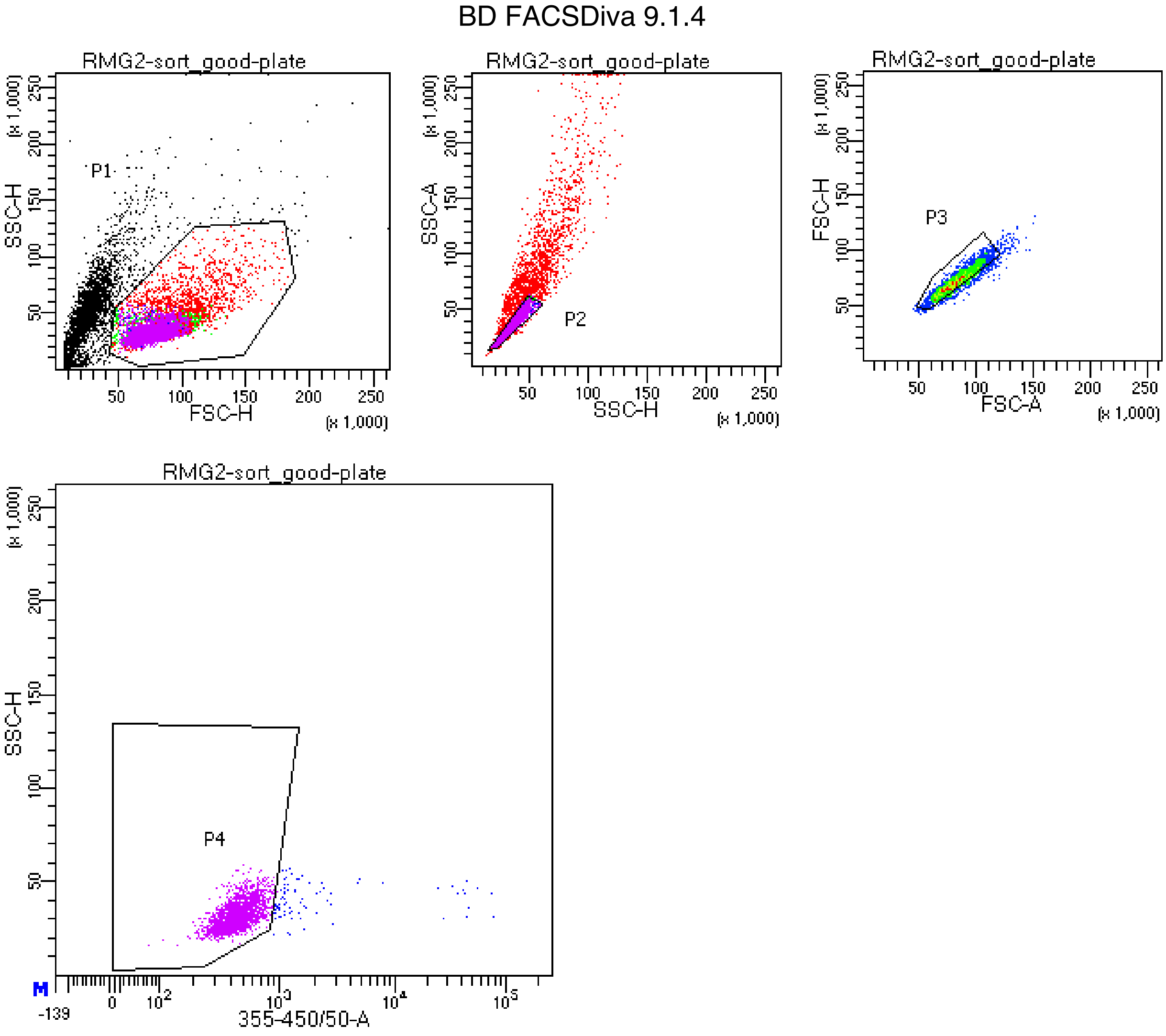


Tube: sort_good-plate

| Population | #Events | %Parent | %Total |
|---|---|---|---|
| All Events | 7,659 | #### | 100.0 |
| P1 | 4,077 | 53.2 | 53.2 |
| P2 | 2,314 | 56.8 | 30.2 |
| P3 | 2,032 | 87.8 | 26.5 |
| P4 | 1,948 | 95.9 | 25.4 |

**Supplemental Figure 17. Representative RMG-II (RMG-2) flow cytometry gating scheme for STORM-seq libraries.** Gating strategy gating on cells (P1), single cells (P2), single cells (P3), and viablity (P4 - DAPI).

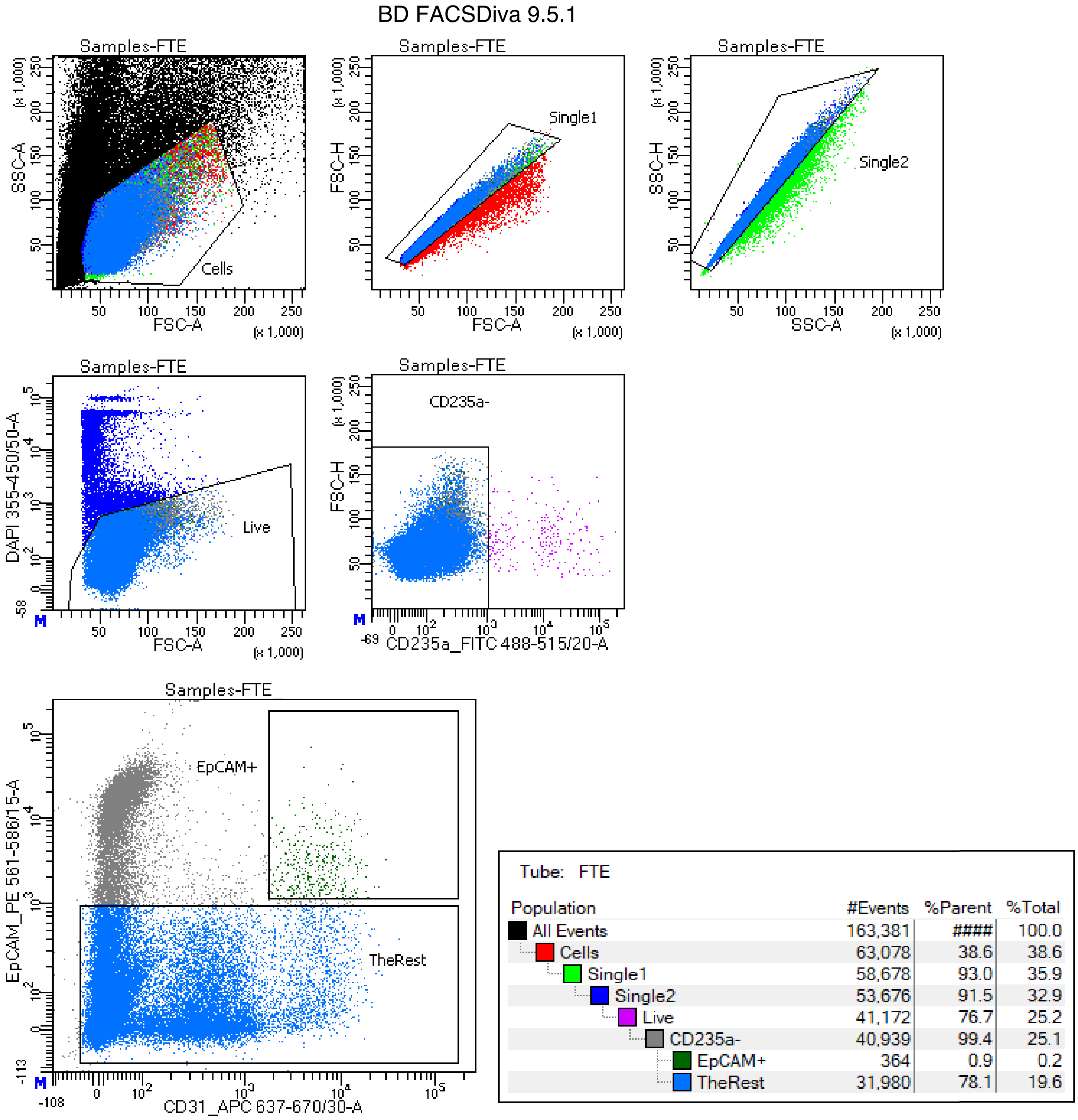


| Population | #Events | %Parent | %Total |
|---|---|---|---|
| All Events | 163,381 | #### | 100.0 |
| Cells | 63,078 | 38.6 | 38.6 |
| Single1 | 58,678 | 93.0 | 35.9 |
| Single2 | 53,676 | 91.5 | 32.9 |
| Live | 41,172 | 76.7 | 25.2 |
| CD235a- | 40,939 | 99.4 | 25.1 |
| EpCAM+ | 364 | 0.9 | 0.2 |
| TheRest | 31,980 | 78.1 | 19.6 |

**Supplemental Figure 18. Representative STORM-seq fallopian tube epithelium gating scheme for STORM-seq libraries.** Gating strategy gating on cells (Cells), single cells (Single1), single cells (Single2), viablity (Live - DAPI), non-RBC (CD235a-), and epithelial cells (EpCAM+).

a

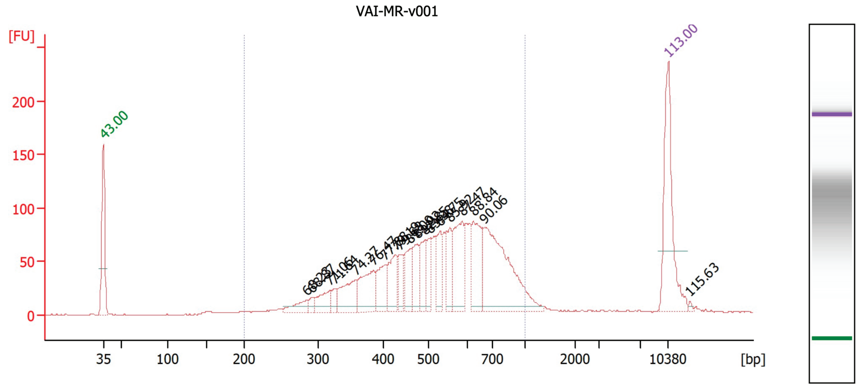


b

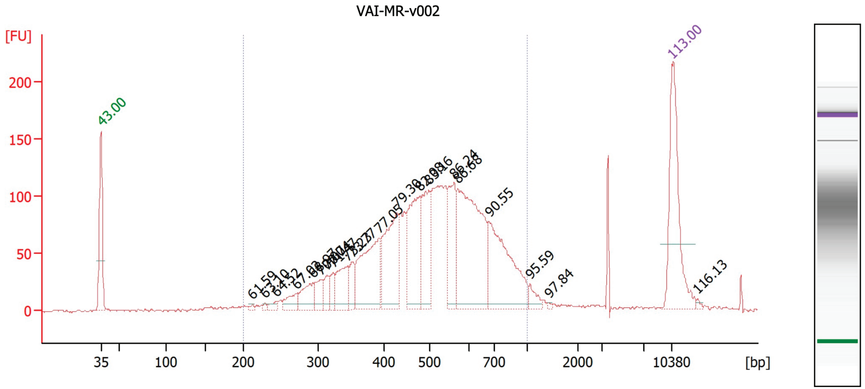


**Region table for sample 10 : VAI-MR-v001**

| From [bp] | To [bp] | Corr. Area | % of Total | Average Size [bp] | Size distribution in CV [%] | Conc. [pg/µl] | Molarity [pmol/l] |
|---|---|---|---|---|---|---|---|
| 200 | 1,000 | 1,778.4 | 95 | 540 | 29.2 | 758.09 | 2,424.8 |

**Region table for sample 11 : VAI-MR-v002**

| From [bp] | To [bp] | Corr. Area | % of Total | Average Size [bp] | Size distribution in CV [%] | Conc. [pg/µl] | Molarity [pmol/l] |
|---|---|---|---|---|---|---|---|
| 200 | 1,000 | 2,346.1 | 94 | 515 | 29.0 | 968.45 | 3,227.4 |

**Supplemental Figure 19. VASA-seq library bioanalyzer traces for plates 1 and 2 of passage matched HEK293T and K-562.** a) Plate 1 (384-well plate) of passage matched HEK293T and K-562 VASA-seq libraries generated by Single Cell Discoveries. b) Plate 2 (384-well plate) of passage matched HEK293T and K-562 VASA-seq libraries generated by Single Cell Discoveries.
